# Predicting mRNA abundance directly from genomic sequence using deep convolutional neural networks

**DOI:** 10.1101/416685

**Authors:** Vikram Agarwal, Jay Shendure

**Affiliations:** Department of Genome Sciences, University of Washington, Seattle, WA 98195, USA; Howard Hughes Medical Institute, Seattle, WA 98195, USA; Brotman Baty Institute for Precision Medicine, University of Washington, Seattle, WA, USA

**Author notes:** Correspondence to Vikram Agarwal and Jay Shendure.

## Abstract

Algorithms that accurately predict gene structure from primary sequence alone were transformative for annotating the human genome. Can we also predict the *expression levels* of genes based solely on genome sequence? Here we sought to apply deep convolutional neural networks towards this goal. Surprisingly, a model that includes only promoter sequences and features associated with mRNA stability explains 59% and 71% of variation in steady-state mRNA levels in human and mouse, respectively. This model, which we call Xpresso, more than doubles the accuracy of alternative sequence-based models, and isolates rules as predictive as models relying on ChIP-seq data. Xpresso recapitulates genome-wide patterns of transcriptional activity and predicts the influence of enhancers, heterochromatic domains, and microRNAs. Model interpretation reveals that promoter-proximal CpG dinucleotides strongly predict transcriptional activity. Looking forward, we propose the accurate prediction of cell type-specific gene expression based solely on primary sequence as a grand challenge for the field.

## INTRODUCTION

Cellular function is governed in large part by the repertoire of proteins present and their relative abundances. Initial attempts to model the gene regulatory forces specifying the mammalian proteome posited a major role for translational regulation, implying that mRNA levels might be more poorly predictive of protein abundance than is often assumed^1^. However, subsequent reanalyses of those data have shown that as much as 84% of variation in protein levels can be explained by mRNA levels, with transcription rates contributing 73%, and mRNA degradation rates contributing 11%^2^. This work reinforces that view that steady-state protein abundances are highly predictable as a function of mRNA levels^3^.

Although quantitative models that predict protein levels from mRNA levels are available^4^, we lack models that can accurately predict mRNA levels. Steady-state mRNA abundance is governed by the rates of transcription and mRNA decay. For each gene, a multitude of regulatory mechanisms are carefully integrated to tune these rates and thus specify the concentrations of the corresponding mRNAs that cells of each type will produce. Key mechanisms include: i) the recruitment of an assortment of transcription factors (TFs) to a gene’s promoter region, ii) epigenetic silencing, as frequently demarcated by Polycomb-repressed domains associated with H3K27me3 histone marks^5^, iii) the activation of genes by enhancers, stretch enhancers^6^, and super-enhancers^7^, frequently associated with H3K27ac histone marks and the Mediator complex, iv) the degradation of mRNA through microRNA-mediated targeting^8^ and PUF family proteins^9^, and v) the stabilization of mRNA through the recruitment of HuR^10^. Jointly modeling these diverse aspects of gene regulation within a quantitative framework has the potential to shed light on their relative importance, to elucidate their mechanistic underpinnings, and to uncover new modes of gene regulation.

Previous attempts to model transcription and/or mRNA decay can be broadly split into those based on correlative biochemical measurements and those based on primary sequence. In the former category, there have been several attempts to model the relationship between TF binding, histone marks, and/or chromatin accessibility and gene expression (*e.g.* predicting the expression levels of genes based on ChIP-seq and/or DNase I hypersensitivity data)^11–17^. While such models can clarify the relationships between heterogeneous, experimentally-derived biochemical marks and transcription rates, their ability to deliver mechanistic insights is limited. For example, for models relying on histone marks, the temporal deposition of such marks might follow, rather than precede, the events initiating transcription, in which case the histone marks reinforce or maintain, rather than specify, a transcriptional program. For models relying on measurements of TF binding, a substantial fraction of ChIP-seq peaks lack the expected DNA binding motif, and potentially reflect artifactual binding signals originating from the predisposition of ChIP to pull down highly transcribed regions or regions of open chromatin^18–20^.

In the latter category, there have also been a few attempts to model transcript levels or mRNA decay rates based solely on primary sequence. For example, a model of the spatial positioning of *in silico* predicted TF binding sites relative to transcriptional start sites (TSSs) was able to explain 8-28% of variability in gene expression^14^. Models based on simple features including the GC content and lengths of different functional regions (e.g., the 5′ UTR, ORF, introns, and 3′ UTR) and ORF exon junction density^21,22^ explain as much as 40% of the variability in mRNA half-lives, which are in turn estimated to explain 6-15% of the variability in steady-state mRNA levels in mammalian cells^1,2,22^. However, the vast majority of variation in in steady-state mRNA levels has yet to be explained by sequence-based models.

To what extent is gene expression predictable directly from genome sequence? Relevant to this question, a recent study relying on a massively parallel reporter assay (MPRA) demonstrated that the transcriptional activities associated with isolated promoters can explain a majority (~54%) of endogenous promoter activity23. This result establishes a clear mechanistic link between the primary sequence of promoters and variability in gene expression levels. This in turn implies that there may exist a mathematical function which, if properly parameterized, could accurately predict mRNA expression levels based upon nothing more than genomic sequence. However, it remains unknown whether such a function is “learnable” given limited training data and highly incomplete domain-specific knowledge of the parameters governing gene regulation [e.g., biochemical parameters describing the affinity of TFs to their cognate motifs (K_d_), constants describing the rates of TF binding and unbinding (K_on_ and K_off_, respectively), potential cooperative effects from the combinatorial binding of TFs (as measured by Hill coefficients), the distance dependencies between TF binding relative to the TSS and RNA Polymerase II recruitment, and competition for binding between TFs and histones^24^, etc.].

Methods based upon deep learning are providing unprecedented opportunities to automatically learn relationships among heterogeneous data types in the context of incomplete biological knowledge^25^. Such methods often employ deep neural networks, in which multiple layers are employed hierarchically to parametrize a model which transforms a given input into a specified output. For example, deep convolutional neural networks have been used to predict the binding preferences of RNA and DNA binding proteins^26^, the impact of noncoding variants on the chromatin landscape^27^, the chromatin accessibility of a cell type from DNA sequence^28^, and genome-wide epigenetic measurements of a cell type from DNA sequence^29^.

The application of deep learning to model the various regulatory processes governing gene expression in a unified framework has great potential, and could enable the discovery of fundamental relationships between primary DNA sequence and steady-state mRNA levels that have heretofore remained elusive. To this end, we introduce Xpresso, a deep convolutional neural network that jointly models promoter sequences and features associated with mRNA stability in order to predict steady-state mRNA levels.

## RESULTS

### An optimized deep learning model to predict mRNA expression levels

We aspired to train a quantitative model utilizing nothing more than genomic sequence to predict mRNA expression levels. To simplify the prediction problem, we first evaluated the correlation structure of 56 human cell types in which mRNA expression levels had been collected and normalized by the Epigenomics Roadmap Consortium^30^. An evaluation of the pairwise Spearman correlations of mRNA expression levels among cell types revealed that most cell types were highly correlated, exhibiting an average correlation of ~0.78 between any pair of cell types (**Figure S1**). This suggests that the relative abundances of mRNA species are largely consistent among all cell types, justifying the initial development of a cell type-agnostic model for predicting median mRNA expression levels. We observed that median mRNA levels for chrY genes were highly variable due to the sex chromosome differences among cell types. Histone mRNAs were also undersampled and measured inaccurately because of the dependency of the underlying RNA-seq protocols on poly(A)-tails, which histones lack. We therefore excluded chrY and histone genes from our analyses.

Given the lack of deep neural networks available in this prediction context, we sought to perform a search of hyperparameters defining the architecture of a neural network that could more optimally predict gene expression levels while jointly modeling both promoter sequences and sequence-based features correlated with mRNA decay (**Figure 1A**). During this search, we varied several key hyperparameters defining the deep neural network, including the sequence window of the promoter to consider, the batch size for training, the number of convolutional and max pooling layers in which to feed the promoter sequence, and the number of densely connected layers preceding the final output neuron (**Table 1**). The mRNA decay features, which included the GC content and lengths of different functional regions (e.g., the 5′ UTR, ORF, introns, and 3′ UTR) and ORF exon junction density^21,22^, were not varied.

**Table 1.** Search space and hyperparameters discovered. Within brackets, the upper and lower range are listed if the variable is either discrete or continuous, and all possible values are listed if the variable is categorical. If the variable is discrete, the step size is provided. Beneath each underlined hyperparameter are nested hyperparameters that are searched. For example, if three [Convolutional → Max pooling] layers are selected, each of these three layers possesses four additional types of hyperparameters to search amongst. The values for consecutive layers that were ultimately selected in the final models are separated by a ‘/’.

| Hyperparameter | Range | Step size<br>(if discrete) | Best identified<br>manually | Best identified<br>from search |
| --- | --- | --- | --- | --- |
| <u>Batch size</u> | $2^{[5,7]}$ | 1 | 64 | 128 |
| <u>Upstream distance from TSS</u> | [-10000,0] | 500 | -1500 | -7000 |
| <u>Downstream distance from TSS</u> | [0, 10000] | 500 | 1500 | 3500 |
| <u>[Convolutional → Max pooling] layer(s)</u> | [One, Two,<br>Three, Four] |  | Two | Two |
| Number of convolutional filters | $2^{[4,7]}$ | 1 | 64 / 64 | 128 / 32 |
| Convolution filter length | [1, 10] | 1 | 5 / 5 | 6 / 9 |
| Convolution dilation rate | [1, 4] | 1 | 1 / 1 | 1 / 1 |
| Max pooling pool size/stride | [5, 100] | 5 | 10 / 20 | 30 / 10 |
| <u>Densely connected layer(s)</u> | [One, Two] |  | One | Two |
| Number of neurons in layer | $2^{[1,8]}$ | 1 | 100 | 64 / 2 |
| Dropout probability | [0,1] |  | 0.5 | 0.00099 /<br>0.01546 |

**Figure 1.**
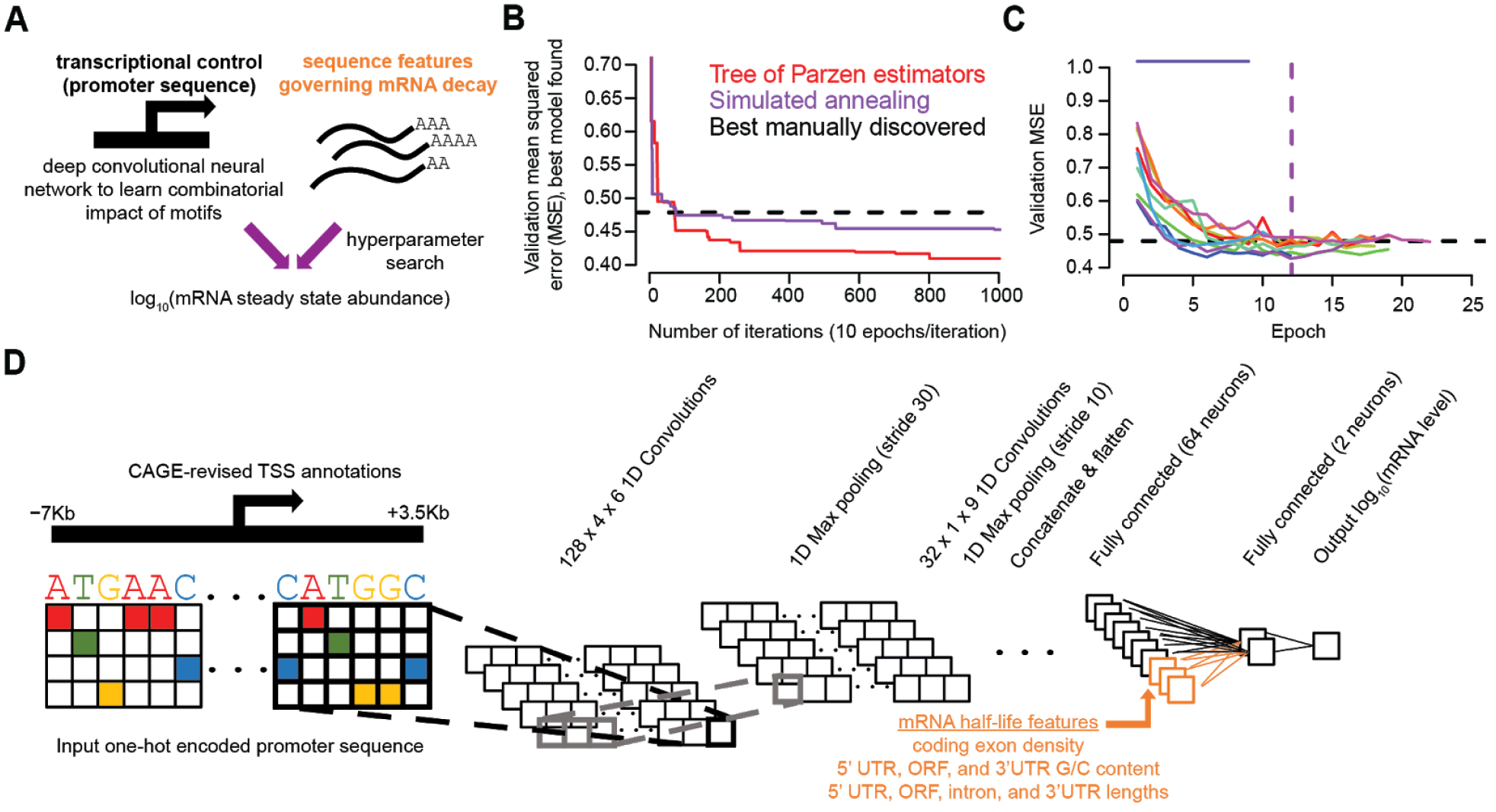
Overview of optimization strategy for deep learning-based model training scheme. **A)** Overview of a predictive model of mRNA steady state abundance that integrates information from sequences at the promoter region of a gene and annotation-based sequence features associated with mRNA decay. **B)** Validation error associated with the best model found at each iteration during the search for an optimal set of hyperparameters to predict median mRNA abundance across tissues. Compared are two strategies (i.e., Tree of Parzen estimators and Simulated annealing) against the performance of the best manually discovered deep learning architecture. **C)** Performance of ten independent trials given the optimal architecture discovered in (B). Nine of ten trials achieve convergence, with the vertical purple dashed line indicating the best model achieved was at the twelfth epoch, and the horizontal dashed line indicating the best manually discovered model as shown in (B). **D)** Best deep learning architecture discovered during the hyperparameter search in (B) corresponds to an architecture with two sequential convolutional and max pooling layers followed by two sequential fully connected layers.

We applied two strategies which have shown promise in the context of hyperparameter searches: the simulated annealing (SA) and the Tree of Parzen estimators (TPE)^31^, optimization strategies that iteratively randomly sample sets of hyperparameters before converging on a set that minimizes the error rate on a validation set. We compared the performance of these methods to the best deep learning architecture defined manually, which was guided by prior knowledge that information governing transcription rate is most likely localized to sequence elements within ±1500bp promoter around a TSS, and further inspired by a deep learning architecture previously used to predict regions of chromatin accessibility from DNA sequence (**Table 1**)^28^. We observed that each optimization strategy progressively discovered better sets of hyperparameters, with the most of the improvements occurring within 200 iterations (**Figure 1B**). The TPE method outperformed the SA strategy, identifying a model whose validation mean squared error (MSE) was 0.401, substantially better than the MSE of 0.479 derived from the best manually defined model.

Given the stochastic nature of training deep learning models, which depend on both the initial parameter configurations and trajectory of the optimization procedure’s search, we devised a strategy to train ten independent trials using the best deep learning architecture specified by the hyperparameters discovered using TPE method. Measuring the validation MSE as a function of the training epoch for these ten trials, we observed one trial that did not converge and nine that converged to similar MSE values (**Figure 1C**). For our final model, we selected the parameters derived from the specific trial and epoch that minimized the validation MSE. All following results of this study report the performance derived from the best of ten trained models.

Our final model, which considered the region 7Kb upstream of the TSS to 3.5Kb downstream of the TSS, was comprised of two sequential convolutional and max pooling layers followed by two sequential fully connected layers preceding the output neuron (**Table 1**, **Figure 1D**, and **Figure S2A**), and consisted of 112,485 parameters in total (**Figure S2A**). While this large 10.5Kb sequence window was the best discovered, an evaluation of suboptimal hyperparameters suggested this expansive upstream region was not critical for good performance, as an alternative model spanning the region 1.5Kb upstream to 7.5Kb downstream of the TSS obtained a similar validation MSE (**Figure S2B**). Furthermore, our manually defined architecture, spanning the region 1.5Kb upstream to 1.5Kb downstream of the TSS achieved an r^2^ of 0.53, only 6% worse than the model discovered by TPE, indicating that a localized region around the core promoter region captured the majority of learnable information, with only a modest additional contribution gained from the consideration of surrounding regions.

To evaluate the relationship between the number of genes in the training set and the performance of the model, we subsampled the training set and evaluated the MSE and r^2^ on the validation and test set, respectively. We found that the greatest gain in performance occurred between 4,000 and 6,000 training examples, with performance on the validation and test sets improving only modestly between 6,000 and 16,000 training examples (**Figure 2A**). The best model trained on the full training set achieved an r^2^ of 0.59.

**Figure 2.**
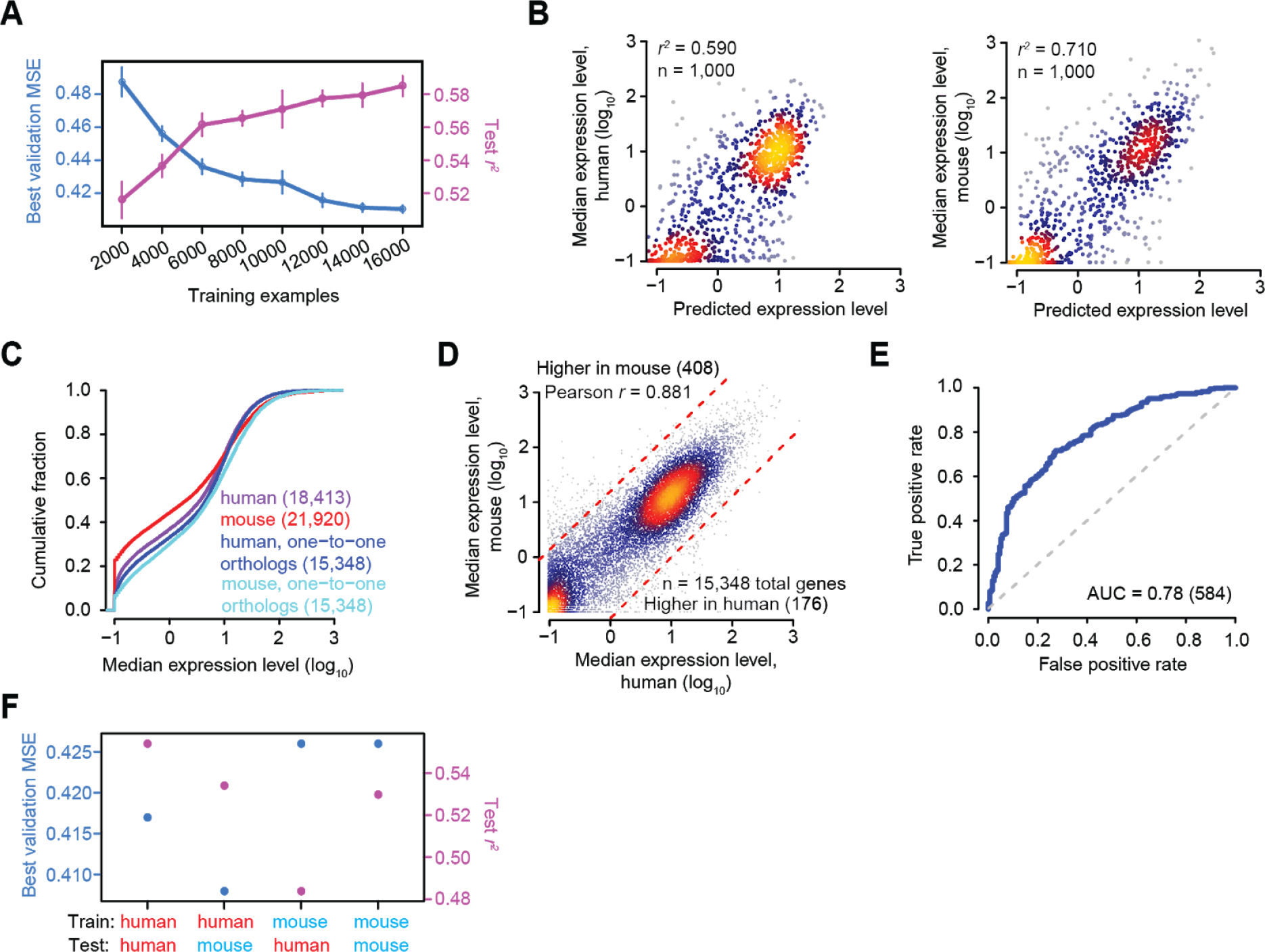
Predictive models explain variability in gene expression in the human and mouse. **A)** Impact of downsampling the training set size on the validation error and performance on the test set [as measured by variance in mRNA expression levels explained (r^2^)]. To generate the mean and 95% confidence intervals displayed, the training set was downsampled (without replacement) into ten batches. For each batch, the performance for the best of ten models, defined as the model with the minimal validation error, was computed. **B)** Scatter plots indicating the performance of human (left panel) and mouse (right panel) models on held-out test sets. Regions are colored according to the density of data from light blue (low density) to yellow (high density). **C)** Cumulative distributions of median mRNA expression levels among tissues for all annotated human and mouse genes as well as those corresponding to the subset of one-to-one human-to-mouse orthologous genes. **D)** Scatter plot indicating the relationship between median mRNA expression levels in human and mouse for one-to-one orthologs. The dotted red lines correspond to the threshold utilized to call species-specific genes, corresponding to a 10-fold change in expression in one species relative to the other. The number of species-specific genes surpassing this threshold is indicated in parentheses. Regions are otherwise colored as in panel (B). **E)** Performance of a classifier that utilizes the difference between cross-validated species-specific predictions to distinguish mRNAs whose expression is strongly enriched in the mouse or human. Shown is a Receiver Operating Characteristic (ROC) curve showing the relationship between False Positive Rate and True Positive Rate at varying thresholds of the predicted expression difference between cell types, with the grey dotted line indicating the expected curve for a classifier performing at random chance. Also shown is the Area Under the Curve (AUC) to quantify performance of the classifier. **F)** Impact of training and testing model performance within and across mammalian species using a test set matched for the same set of one-to-one orthologs. Shown is the performance for the best of ten models acquired for the full training set derived from each species.

### Performance of predictive models in human and mouse

We next sought to compare the generality and performance of our method across mammalian species. We focused on 18,377 and 21,856 genes in human and mouse, respectively, for which we could match promoter sequences and gene expression levels, and held out 1,000 genes in each species as a test set. With the aforementioned model architecture, we trained and tested a model for predicting the median mRNA expression levels of mouse genes based on mouse genomic sequence, achieving an r^2^ of 0.71, substantially higher than the r^2^ of 0.59 achieved with the human model (**Figure 2B**). Interested in explaining this 12% discrepancy, we compared the distribution of median mRNA expression levels between species. While over 20% of mouse genes were non-expressed (**Figure 2C**, displayed on the x-axis as −1 due to the addition of a pseudocount of 0.1 prior to log-transforming the RPKM values), fewer than 10% of human genes were non-expressed. In contrast, evaluating the subset of 15,348 one-to-one orthologs in human and mouse revealed a similar proportion of non-expressed genes (**Figure 2C**). These one-to-one orthologs showed strikingly concordant median expression levels (**Figure 2D**), indicating that gene expression levels of these two species have remained heavily conserved since the divergence of a common mammalian ancestor ~80 million years ago.

Of note, a subset of genes did exhibit substantial differences in expression between mouse and human. We identified a cohort of 584 mRNAs enriched by at least 10-fold in one species; of these, 176 were relatively up-regulated in the mouse and 408 in human (**Figure 2D**). To what extent can these species-specific differences be explained on the basis of differences in promoter sequence? We next compared our mouse and human Xpresso models to evaluate how well these models could discriminate species-specific mRNAs. A binary classifier based upon the difference in predictions from each species could indeed correctly discriminate these species-specific mRNAs better than chance expectation (AUC = 0.78, **Figure 2E**). This observation beckoned the question of whether the sequence–expression relationships learned in each species were of a similar nature.

To test whether the regulatory rules learned by each model could generalize across species, we retrained human and mouse-specific models using training and validation sets that were matched to have the same groups of one-to-one orthologs. We then tested the performance of these models on a held-out group of one-to-one orthologs in either the same species or the opposite species. While the best performing models were trained on the same species, each model performed only marginally worse when tested on the opposite species (i.e., within a 6% decrease in r^2^) (**Figure 2F**), demonstrating that the regulatory principles learned by the deep learning model generalize across the mammalian phylogeny. The similar r^2^ values between human and mouse obtained when restricting the analyses to one-to-one orthologs implies that the greater r^2^ previously observed in the mouse relative to human (**Figure 2B**) was due to the mouse model simply discriminating expressed genes from the mouse’s larger fraction of non-expressed genes. To directly test this possibility, we downsampled the proportion of non-expressed mouse genes to match that of the human, and re-trained a mouse-specific model. The r^2^ decreased only modestly to 0.65 on a held-out test set, suggesting that the differential proportion of non-expressed genes only partially explains the improved performance in the mouse, with other factors such as data quality potentially playing a role.

### Cell type-specific models implicate a diversity of gene regulatory mechanisms

Given the generality of our deep learning framework, we next sought to build cell type-specific models. With the same hyperparameters, we trained new models to predict the expression levels of all protein-coding genes for human myelogenous leukemia cells (K562), human lymphoblastoid cells (GM12878), and mouse embryonic stem cells (mESCs) (**Supplementary Table 1**), cell types for which abundant functional data is available. To avoid overfitting, we developed a 10-fold cross-validation-based procedure to ensure that the prediction for any given gene resulted from its being part of a held-out sample (i.e., it was not part of the training set from which the model that predicted its expression level was built). From the residuals (i.e., the difference between actual expression levels and our predictions based solely on promoter sequence and mRNA decay rate features), we investigated whether we could observe the influence of additional gene regulatory mechanisms that were not initially considered, or incompletely accounted for, in the Xpresso model.

We first evaluated K562 cells, finding that our cell type-specific predictions correlated with observed K562 expression levels with an r^2^ of 0.51 (**Figure 3A**). This ~8% decrease in performance relative to cell type-agnostic predictions (**Figure 2B**) suggested that the expression levels of tissue-specific genes may be harder to predict than those of housekeeping genes, because such genes might be under the control of gene regulatory mechanisms not considered by our model. One obvious such mechanism involves enhancers, *cis*-acting regulatory elements that may be located hundreds of kilobases away from a TSS. For example, in K562 cells, distal enhancers have been implicated as modulating the expression of most genes of the α-globin and β-globin loci, *GATA1*, *MYC*, and others^32–34^. Reasoning that such genes should be consistently underestimated by our predictions, we plotted the distribution of their residuals (**Figure 3A**). Indeed, all of these genes were expressed much more highly than our predictions in K562 cells, reinforcing the notion that such genes are activated by regulatory mechanisms beyond promoters. Among the biggest outliers were the β-globin genes, which in some cases were expressed over four orders of magnitude more highly than predicted (**Figure 3A**). Collectively, the residuals corresponding to all of these known enhancer-driven genes were heavily biased towards positive values relative to all other genes (*P* < 10^−9^, one-sided Kolmogorov-Smirnov (K-S) test, **Figure S3A**), confirming that the model consistently under-predicted their expression levels.

**Figure 3.**
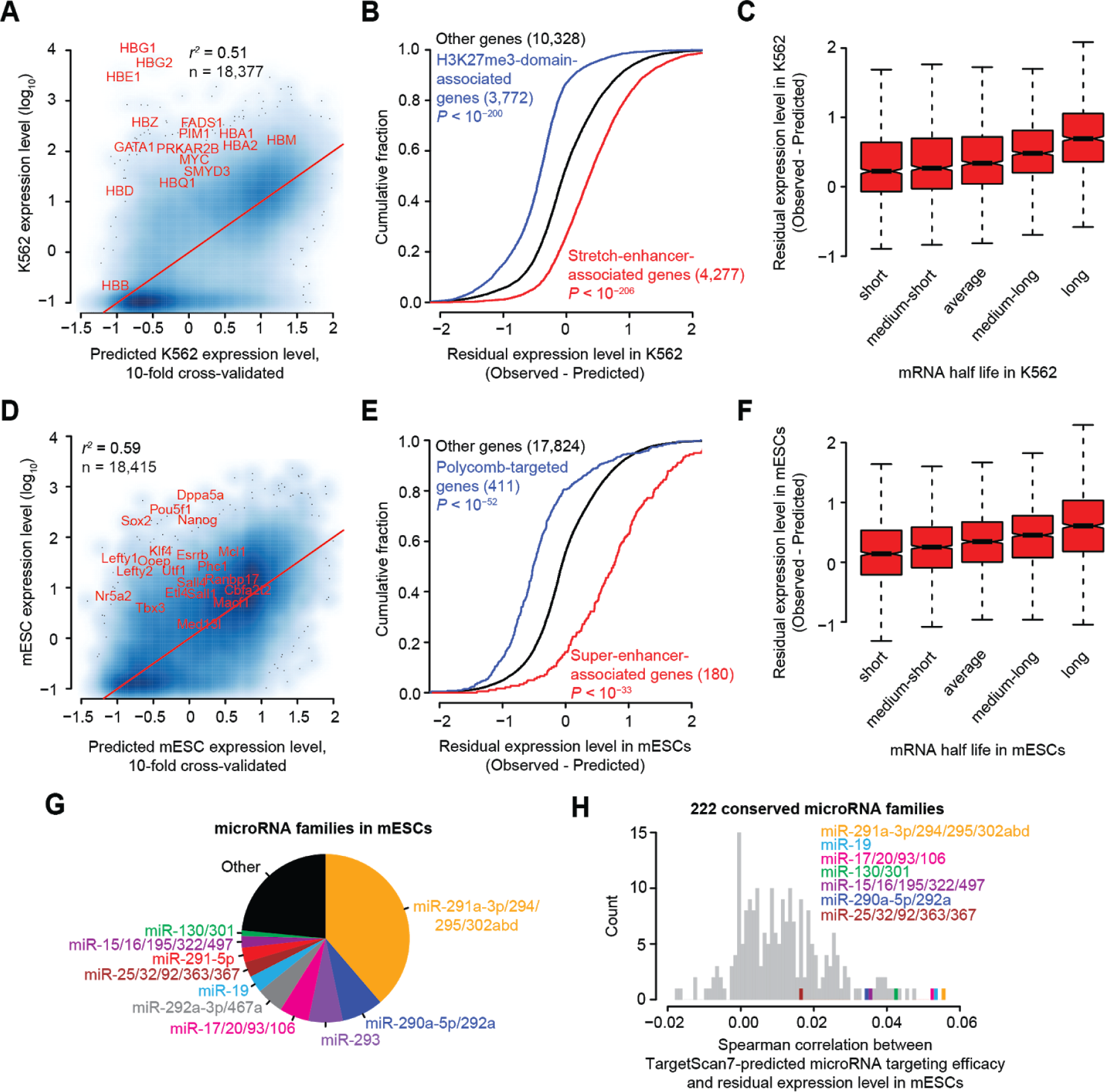
A diversity of gene regulatory mechanisms are associated with the residuals of cell type-specific models. **A)** Relationship between 10-fold cross-validated predictions and actual mRNA expression levels in K562 cells. Regions are colored according to the density of data from light blue (low density) to dark blue (high density). Labeled in red are the globin genes alongside others implicated as genes activated by strong enhancers in K562 cells. Gene names were moved slightly to enhance readability. **B)** Cumulative distributions of residuals corresponding to all stretch-enhancer-associated genes, H3K27me3-domain-associated genes, and control genes not associated with either. Similarity of the distributions to that of the set of controls was tested (one-sided Kolmogorov–Smirnov [K–S] test, *P* value); the number of mRNAs analyzed in each category is listed in parentheses. **C)** Boxplots showing the relationship between mRNA half-life and residuals in K562 cells; indicated is the median residual value (bar), 25th and 75th percentiles (box), and the minimum of either 1.5 times the interquartile range or the most extreme data point (whiskers). half-life measurements were measured using TimeLapse-seq^36^ in K562 cells (n = 5,007 genes), and partitioned into 5 equally-sized bins spanning the range of half-life values. **D)** This panel mirrors that shown in (A), except that it highlights genes associated with known enhancers and super-enhancers in mouse embryonic stem cells (mESCs). **E)** This panel mirrors that shown in (B), except that it compares genes associated with super-enhancers and Polycomb-repressed domains in mESCs. **F)** This panel mirrors that shown in (C), except that half-life measurements were measured using SLAM-seq^39^ in mESCs (n = 6,266 genes). **G)** Pie chart indicating the relative proportions of microRNA families expressed in mESCs. Colored are the top 10 most abundant miRNA families. **H)** Histogram of the Spearman correlation values between TargetScan7-predicted microRNA targeting efficacy^8^ and residual expression level in mESCs, for 222 miRNA families conserved among mammals. Highlighted are the subset of 7 highly abundant miRNA families colored in panel (G) that are also conserved among mammals.

To more systematically identify genes activated or silenced by non-promoter mechanisms, we developed a method to predict them genome-wide. Given that enhancers are frequently associated with large domains of H3K27Ac activity, and that silenced genes are frequently associated with heterochromatic domains marked by H3K27me3, we examined genome-wide chromatin state annotations based upon diHMM, a method that annotates such domains genome-wide in K562 and GM12878 cells using histone marks^35^. Although a subset of H3K27Ac-associated domains were originally called “super-enhancers”^35^, we find it more appropriate to refer to them as “stretch enhancers”, which are more loosely defined as clusters of enhancers spanning ≥3Kb^6^. In total, we identified 4,277 genes that overlap stretch enhancers, and 3,772 genes that overlap with H3K27me3-domains, ignoring those that happen to overlap with both. Consistent with our expectation, these collections of genes were significantly associated with predominantly positive and negative residuals, respectively, relative to the background distribution of other genes (*P* < 10^−200^, one-sided K-S test, **Figure 3B**). The same was true for GM12878 cells (**Figure S3B**), reinforcing the generality of this finding across cell types.

We next examined whether our residuals could be further explained by post-transcriptional gene regulatory mechanisms. Highly reproducible mRNA half-life estimates for 5,007 genes in K562 cells were previously measured using TimeLapse-seq^36^. We observed a positive correlation between mRNA half-lives (which were log-transformed) and our residuals (Pearson correlation = 0.28; *P* < 10^−15^). We visualized this trend by splitting the half-lives into five equally-sized bins (**Figure 3C**). These results show that although we included sequence-based features associated with mRNA decay rates in the model, these were insufficient to capture the full contribution of mRNA decay rates to steady-state mRNA levels.

We next turned our attention to mouse ESCs. Key stem cell identity genes include *Pou5f1* (also known as *Oct4*), *Sox2*, *Nanog*, and a host of others whose dependence upon enhancers and super-enhancers has been experimentally validated with CRISPR-deletion experiments^7,37^. Similar to the other cell types, although a mESC-specific model could strongly predict mRNA expression levels in mESCs (r^2^ = 0.59), key stem cell identity genes harbored residuals that were strongly biased towards positive values (**Figure 3D**, **Figure S3C**), confirming that their promoter sequences and mRNA sequence features could not adequately explain their high abundance. Extending this question more systematically to 180 protein-coding genes thought to be governed by super-enhancers in mESCs^7^, we observed a strong enrichment for highly positive residuals in these genes relative to all other genes (*P* < 10^−33^, one-sided K-S test; **Figure 3E**).

In mESCs, genes associated with the Polycomb Repressive Complex (PRC), as delineated by binding to both PRC1 and PRC2 and frequently marked with H3K27me3, are thought to be associated with key developmental regulators, many of which are silenced but poised to be activated upon differentiation^38^. We observed that this group of genes, in contrast to the super-enhancer-associated set, exhibited a strong enrichment in highly negative residuals relative to all other genes (*P* < 10^−52^, one-sided K-S test; **Figure 3E**), consistent with a model in which PRC-targeted genes are actively silenced.

Mirroring our analysis from human cells, we next evaluated the relationship between mRNA half-lives in mESCs and our residuals. We obtained reproducible mRNA half-life estimates for 6,266 genes in mESCs measured using SLAM-seq^39^. Similar to human cell types, residuals were positively correlated with mRNA half-lives (Pearson correlation = 0.24; *P* < 10^−15^; **Figure 3F**).

A distinguishing feature of mESCs relative to K562 cells is that more is known about the post-transcriptional regulatory mechanisms governing mRNA half-life. In particular, microRNAs serve as strong candidates for further inquiry as they are guided by their sequence to bind and repress dozens to hundreds of mRNAs, mediating transcript degradation and thereby shortening an mRNA’s half-life. The miR-290-295 locus, essential for embryonic survival, encompasses the most highly abundant miRNAs in mESCs. Under the control of a super-enhancer, the members of this miRNA cluster are expressed in a highly cell type-specific manner and are thought to operate as key post-transcriptional regulators in ESCs^7^. We asked whether we could use our residuals to infer the endogenous regulatory roles of abundant miRNAs in mESCs. Defining a miRNA family as any miRNA sharing an identical seed sequence (as indicated by positions 2-8 relative to the miRNA 5′ end^8^), we used existing small RNA sequencing data from mESCs^40^ (GSE76288) to quantify miRNA family abundances for the top 10 miRNA families. In addition to the miR-290-295 family, we detected other highly abundant families including miR-17/20/93/106, miR-19, miR-25/32/92/363/367, miR-15/16/195/332/497, and miR-130/301; these miRNA families collectively comprised over 75% of the total miRNA pool in mESCs (**Figure 3G**).

For each of the 222 miRNA families conserved across the mammalian phylogeny, which includes 7 of the 10 top miRNA families in mESCs, we assessed whether the predicted repression of targets correlated to our residuals. We used the TargetScan7 Cumulative Weighted Context+ Score (CWCS)^8^ to rank predicted conserved targets for the subset of mRNAs expressed in mESCs, assigning a CWCS of zero for non-targets. The miRNA family with the most highly ranked Spearman correlation corresponded to that of the miR-291a-3p/294/295/302abd family (**Figure 3H**), which was also the most highly expressed miRNA family in ES cells. The sign of this correlation was consistent with expectation, as targets with more highly negative CWCSs (corresponding to predicted targets with greater confidence) had negatively shifted residual values. More generally, the 22 miRNA families comprising the highest 10% of Spearman correlations were strongly enriched for the 7 miRNA families highly abundant in mESCs (*P* < 10^−5^, Fisher’s exact test; **Figure 3H**). Our results thus reinforce the finding that highly abundant miRNAs mediate target suppression^41^ while providing an alternative, fully computational method to infer highly active endogenous miRNA families in specific cell types solely from primary sequence and gene expression data.

Collectively, our results from K562, GM12878, and mESCs demonstrate how analyses of Xpresso’s residuals can be used to explore and quantify the influence of diversity gene regulatory mechanisms. The mRNAs with the most positive residuals are highly enriched in genes associated with the activity of enhancers and super-enhancers, while those associated with the most negative residuals are are highly enriched in genes associated with the activity of pathways involved in gene silencing, such as those targeted by Polycomb Repressive Complexes and microRNAs. We anticipate that cell type-specific quantitative models for any arbitrary cell type can serve as a useful hypothesis generation engine for the characterization of active regulatory regions in the genome and key regulators such as miRNAs, including for cell types in which histone ChIP and small RNA sequencing data is limited or unavailable.

### Performance of cell type-specific Xpresso models

To further characterize the ability of Xpresso to learn cell type-specific expression patterns, we evaluated the relative performance of cell type-agnostic and cell type-specific models in predicting cell type-specific mRNA levels. In all three cases considered, models trained on the cell type of origin out-performed those trained on median expression levels by about 3-5% (**Figure 4A**). We next compared our K562 and GM12878 Xpresso models to evaluate how well these models could discriminate cell type-specific mRNAs. We identified a cohort of 1,977 mRNAs enriched by at least 10-fold in each cell type; of these, 890 were relatively up-regulated in K562 cells and 1,087 were up-regulated in GM12878 cells (**Figure 4B**). A binary classifier based upon the difference in predictions from each cell type could discriminate these cell type-specific mRNAs modestly better than chance expectation (AUC = 0.65, **Figure 4C**).

**Figure 4.**
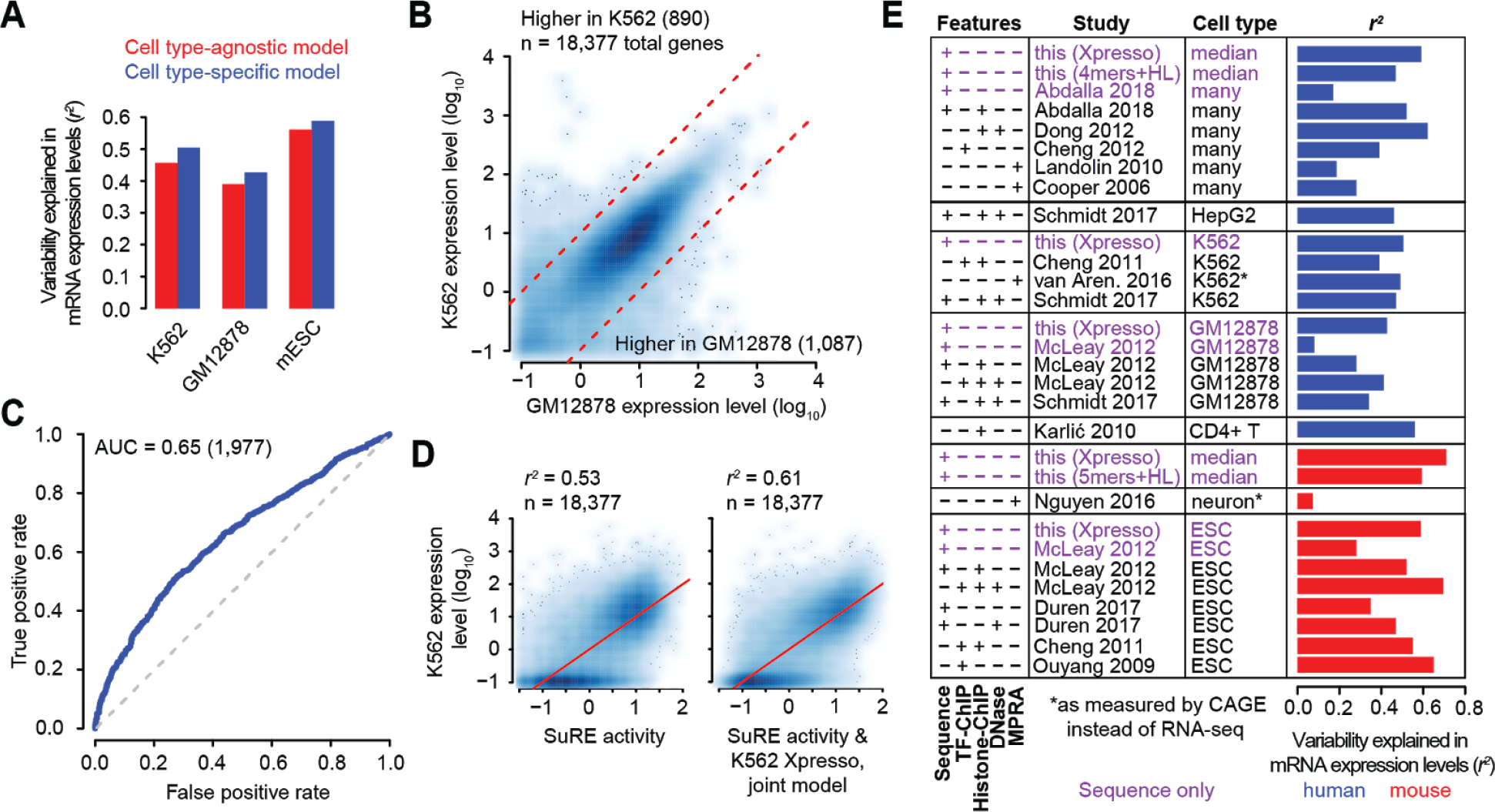
Cell type-specific models are competitive with methods based upon experimental data. **A)** Bar plots comparing the variance explained in mRNA expression levels in each of three cell types (human K562 and GM12878 cells as well as mouse embryonic stem cells), using models trained on median expression levels (i.e., cell type-agnostic model) or on the matched cell type (i.e., cell type-specific model). The r^2^ shown is derived from the entire dataset, using the cross-validated predictions of each strategy. **B)** Relationship between mRNA expression levels in GM12878 and K562 cells. The dotted red lines correspond to the threshold utilized to call cell type-specific genes, corresponding to a 10-fold change in expression in one cell type relative to the other. The number of cell type-specific genes surpassing this threshold is indicated in parentheses. Scatter plots are otherwise colored as in **Figure 3A**. **C)** Performance of a classifier that utilizes the difference between cell type-specific predictions to distinguish mRNAs whose expression is strongly enriched in either GM12878 or K562 cells. Shown is a ROC curve showing the relationship between False Positive Rate and True Positive Rate at varying thresholds of the predicted expression difference between cell types, with the grey dotted line indicating the expected curve for a classifier performing at random chance. Also shown is the AUC to quantify performance of the classifier. **D)** Shown in the left panel is the relationship between SuRE^23^, a Massively Parallel Reporter Assay (MPRA) to measure autonomous promoter activity, and mRNA expression levels in K562 cells. Shown in the right panel is the relationship between a joint SuRE and Xpresso model and mRNA expression levels in K562 cells. Scatter plots are colored as in panel (B). **E)** Comparison of the r^2^ of our sequence-only models to those derived from alternative strategies reported in the literature, often trained using a variety of cell type-matched experimental datasets such as those based upon ChIP of transcription factors or specific histone marks, DNase hypersensitivity measurements, and MPRAs. Methods using nothing more than genomic sequence to predict expression are highlighted in purple, and results in human and mouse are shown in blue and red, respectively.

Next, we sought to estimate the maximum possible performance for predicting gene expression from promoter sequences alone. A recently developed genome-wide MPRA measuring autonomous promoter activity in K562 cells, called Survey of Regulatory Elements or SuRE, linked 200bp to 2Kb regions of the genome to an episomally-encoded reporter in order to measure the transcriptional potential of regulatory sequences^23^. SuRE therefore provides an orthogonal, empirical means of assessing the regulatory information held in promoters, independent of the influence of genomic context and distal regulatory elements. Indeed, we observed that SuRE activity in the ±500bp promoter region around a TSS was highly correlated to mRNA expression levels in K562 cells (r^2^ = 0.53, **Figure 4D**), indicating that just over half of the variation in mRNA expression levels can be explained by regulatory sequences contained within promoters.

The comparable r^2^ achieved by SuRE measurements and the K562-specific Xpresso model in predicting K562 expression levels (r^2^ = 0.53 and 0.51, respectively) provided a unique opportunity to evaluate how well our model captured the experimentally measurable information regarding a promoter’s transcriptional activity. To assess the level of information shared between SuRE measurements and Xpresso predictions, we built a joint model to predict K562 levels. This model achieved an r^2^ of 0.61, 8-10% better than either method alone, suggesting that while both methods captured largely redundant information, each also captured additional information not incorporated in the other (**Figure 4D**). Nevertheless, the relatively modest increase in performance of the joint model over Xpresso alone indicates that Xpresso was able to learn the major sources of sequence-encoded information that explain mRNA expression levels.

### Predictive models perform competitively with models utilizing experimental data

We next evaluated the performance of Xpresso relative to an assortment of baseline and pre-existing models that attempted to predict mRNA levels, both with and without the consideration of mRNA half-life features. For the baseline models, we attempted to predict median expression level using simple *k*-mer counts in the ±1500bp promoter region, the presence of predicted TF binding sites given known motifs available in the JASPAR database, or joint models considering both (**Figure S4A**). These models were trained using simple multiple linear regression and evaluated on the same test set as that utilized in **Figure 2**. Varying the *k*-mer size from *k* = 1 to 6, we found that performance plateaued at *k* = 4 and 5 for human and mouse, respectively, with the greatest gain in performance occurring between *k* = 1 and 2 in both species. Consideration of known TF binding sites in a joint model at best only marginally improved performance, though a model considering these binding sites alone performed as well as a model based on 2-mers.

All of the models benefitted from the additional consideration of half-life features. We evaluated the coefficients associated with a model considering only half-life features to assess the relative contribution of individual features. The features most strongly associated with increased steady-state mRNA abundance in both the human and mouse corresponded to ORF exon density and 5′ UTR GC content, followed by weaker associations to 5′ UTR length and ORF length. In contrast, intron length was negatively associated with mRNA abundance (**Figure S4B**).

Overall, although our baseline models demonstrated that models built upon simple features could perform surprisingly well, our hyperparameter-tuned Xpresso model improved upon these models by 11.2% and 11.7% in human and mouse, respectively (**Figure S4A**). Our 10-fold cross-validation results further verified that Xpresso performed significantly better than the best alternative *k*-mer-based approach in both the human and mouse (*P* < 10^−8^, paired t-test, **Figure S4C**).

Next, we compared our best baseline and Xpresso models to existing models described in the literature, categorizing the types of features used as input in each model into five categories: i) those using nothing more than sequence features, which included our method and two others^14,42^, those using MPRAs to measure promoter activity^23,43–45^, iii) those utilizing the binding signal of transcription factors (TFs) at promoter regions, as measured by ChIP^11,13–15^, iv) those utilizing the signal of histone marks such as H3K4me1, H3K4me3, H3K9me3, H3K27Ac, H3K27me3, and H3K36me3 at promoters and gene bodies, as measured by ChIP^11,12,14,16,17,42^, and v) those utilizing DNase hypersensitivity signal at promoters and nearby enhancers^12,14,16,46^ (**Figure 4E**). Many of these models were trained and tested on cell lines such as K562, GM12878, and mESCs, for which ChIP data is available for a multitude of histone marks and TFs. Thus, we were also able to compare the relative performance of our cell type-specific models for these same cell lines.

The only existing method which attempted to predict expression from sequence features alone14 achieved an r^2^ of 0.08 and 0.28 in GM12878 and mESCs, respectively. In comparison, Xpresso models tested on the same cell types achieved an r^2^ of 0.43 and 0.59, more than doubling the performance of sequence-only models in each (**Figure 4E**). MPRA-based models exhibited a wide diversity of r^2^ values, although the genome-wide MPRA performed in K562 cells^23^ performed comparably to Xpresso. Among all models examined, those utilizing multiple forms of experimental data such as TF ChIP, histone ChIP, and DNase achieved the best r^2^ values of 0.62^12^ and 0.70^14^ in human and mouse, respectively, marginally better than the best Xpresso models in these species for matched cell types.

Thus, we demonstrate that models utilizing nothing more than genomic sequence are capable of explaining mRNA expression levels with as much predictive power as—and often more than—analogous models trained on abundant experimental data. Our models have the advantage that they are simple to train on any arbitrary cell type, including those lacking experimental data such as ChIP and DNase. Furthermore, sequence-only models can further augment the performance of existing models that predict mRNA levels in cell types for which experimental data is already available (**Figure 4A**, **Figure S4**).

### Xpresso predicts genome-wide patterns of transcriptional activity

Convolutional neural networks have recently been used to successfully predict patterns of CAGE activity and histone ChIP signal throughout the genome^29^. To accomplish this feat, a deep convolutional neural network was trained on genome-wide information across entire chromosomes, using 131Kb windows that collectively encompassed ~60% of the human genome^29^. This led us to ask whether it was possible to predict genome-wide CAGE signal using our cell type-agnostic Xpresso model, which was trained, in contrast, on 10.5Kb windows comprising only ~5% of the human genome. As a proof of concept, we fixed the half-life features to equal that of the average gene, and generated promoter activity predictions in 100bp increments along a randomly selected 800Kb region of the human genome that encompassed twenty genes, visualizing whether Xpresso could recapitulate the average pattern of CAGE activity across cell types (**Figure 5**). We observed that the Xpresso predictions faithfully reproduced the pattern of CAGE activity^47,48^ in this region. Peaks of high predicted transcriptional activity frequently corresponded to CpG islands^49^ and promoter regions across multiple cell types as predicted by ChromHMM^50^. Xpresso predicted similar expression signatures for both positive and negative DNA strands, revealing it could not distinguish the strandedness of CAGE signal. Confirming that our results generalize across species, consistent results were observed if Xpresso predictions were generated on the 700Kb syntenic locus of the mouse genome (**Figure S5**).

**Figure 5.**
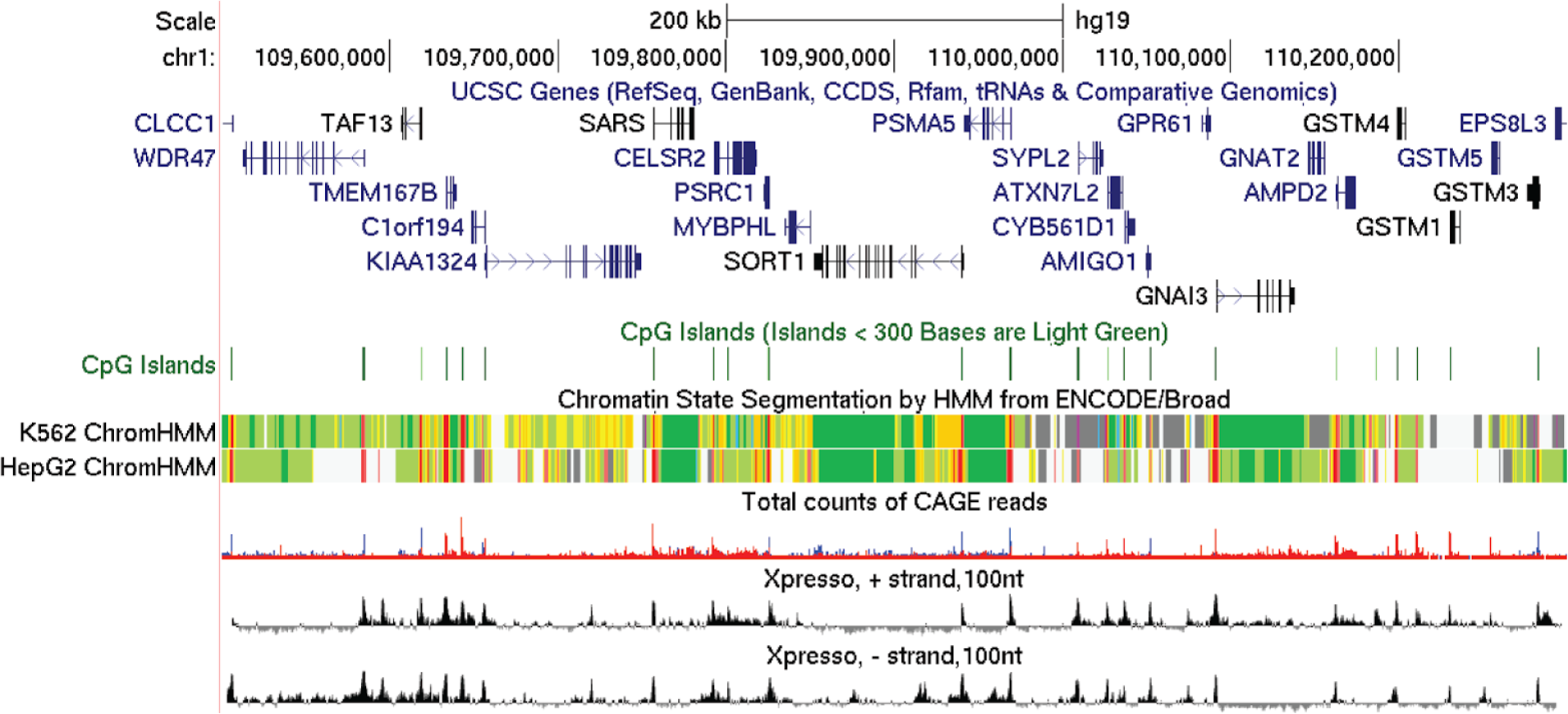
Xpresso predictions recapitulate transcriptional activity across a 800Kb region of human chromosome 1. Shown are the predicted Xpresso-predicted expression levels in 100-nt increments from the plus and minus strands of genomic sequence of the human hg19 genome assembly. Also shown are gene annotations, CpG island calls, ChromHMM genomic state segmentation calls among two cell types (with predicted promoter hidden states colored in red), and genome-wide CAGE signal aggregated among many cell types (with red indicating signal from the positive strand and blue indicating signal from the negative strand). See also **Figure S5** for predictions on the syntenic mouse locus.

### Xpresso automatically identifies the expression level-determining region of the promoter

Interested in ascertaining how our deep learning models could predict cell type-agnostic gene expression levels with high accuracy, we developed a procedure to interpret the dominant features learned and utilized by the mouse and human models. Specifically, we tested four strategies intended to map the regions of the input space of deep learning models with the greatest contribution to the final prediction: i) Gradient * input, ii) Integrated gradients, iii) DeepLIFT (with Rescale rules), and iv) ε-LRP^51^. Each of these methods computed “saliency scores”, which represent a decomposition of the final prediction values into their constituent individual feature importance scores for each nucleotide in the input promoter sequences. We partitioned genes into four groups, including those predicted to be approximately non-expressed and three additional terciles (predicted low, medium, or high expression). We computed mean saliency scores for each of these groups, and then computed the difference in mean scores relative to predicted non-expressed genes for each nucleotide position in the entire input window (i.e., 7Kb upstream of the TSS to 3.5Kb downstream). Averaging our results across our 10-fold cross-validated models, we discovered that the models had automatically learned to consistently rely upon local sequences from the 1Kb sequence centered upon the TSS to predict gene expression (**Figure 6A**, **Figure S6**). Sequences upstream of the TSS contributed more heavily to the predictions than those downstream, with those in the proximal promoter (i.e. within 100bp upstream of the TSS) best discerning genes predicted to have high expression from those predicted to have medium expression (**Figure 6A**). Thus, from only genomic sequence and expression data, the model automatically learned spatial relationships and asymmetries in the relevance of genomic sequence that are consistent with experimental measurements^23^.

**Figure 6.**
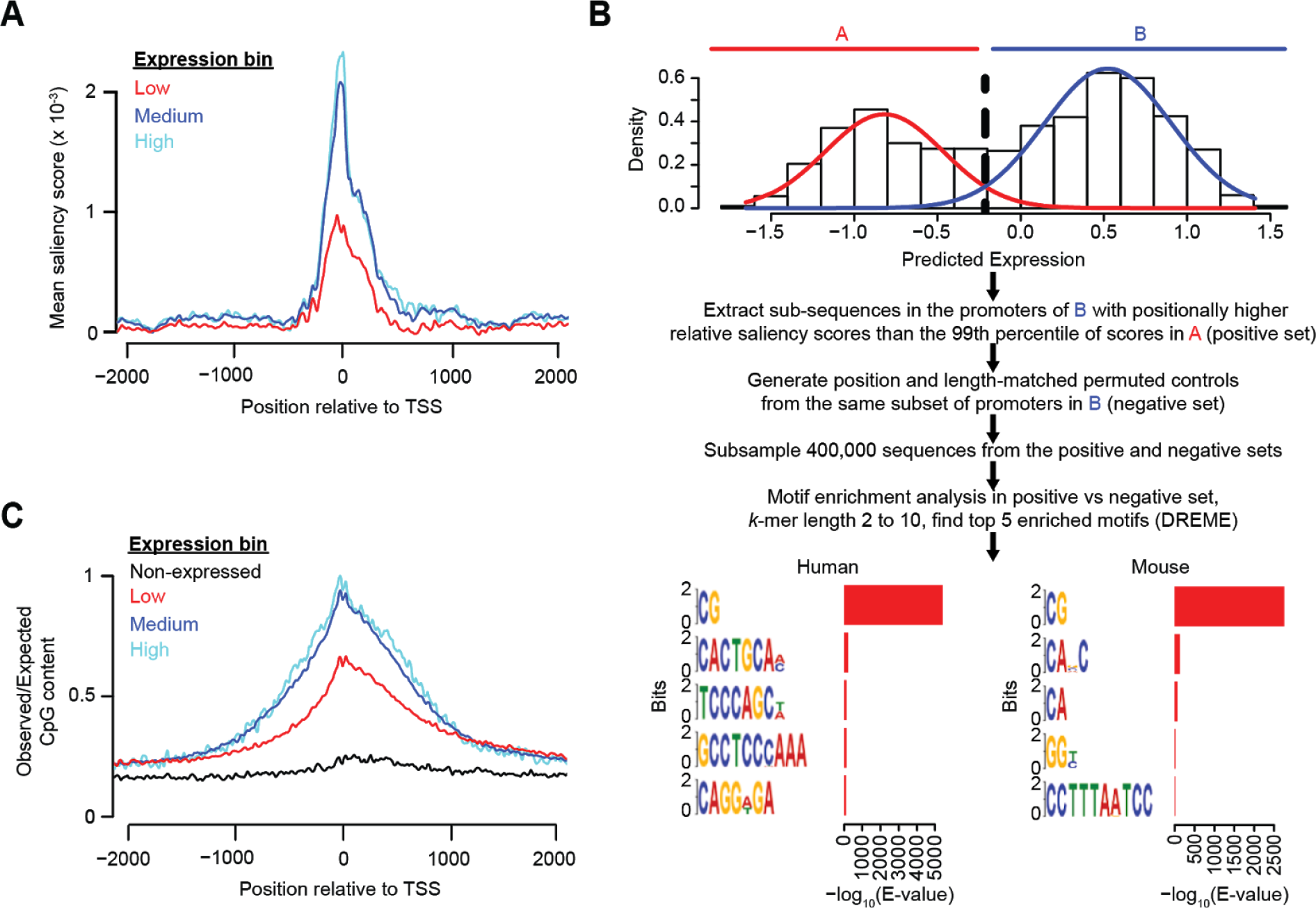
The spatial distribution of CpG dinucleotides in the proximal promoter predicts gene expression levels. **A)** Mean saliency scores relative to predicted non-expressed genes for genes in three terciles of expression bins. Shown are saliency scores from the −2000 to +2000 regions of the input window surrounding the TSS for human promoters as computed using the “Gradient * input” approach. Data has been loess-smoothed at the resolution of 100bp. See also **Figure S6** for the full input region and results using multiple saliency scoring methods in the human and mouse. **B)** Strategy to ascertain enriched *k*-mers in genes predicted to be highly expressed. Shown are the top 5 significantly enriched *k*-mers retrieve from the human and mouse models. **C)** Positional enrichment of CpG dinucleotides relative to chance expectation in human promoters for different gene expression level bins. The expected frequency of CpG dinucleotides is computed as the joint probability of the composite C and G co-occurring based upon their frequency in the corresponding position and gene expression bin. Data has been loess-smoothed at the resolution of 100bp. See also **Figure S7** and **Figure S8** for the results of all dinucleotides in both human and mouse.

Of note, because we used Rectified Linear Units (ReLUs) in our networks, the ε-LRP method resulted in identical results as the Gradient * input method^51^ (Gradient * input: **Figure S6A**; ε-LRP: data not shown). We also observed that the DeepLIFT method led to nearly identical results as the Integrated gradients technique (Integrated gradients: **Figure S6B**; DeepLIFT: data not shown), as observed previously in other contexts^51^. The results in the 10-fold cross-validated mouse model also emulated those of the human (**Figure S6C-D**), indicating that they generalize across species and do not depend upon the specific saliency scoring method used.

### CpG dinucleotides are the dominant signal explaining expression levels

The ability of the model to automatically identify and heavily weight the proximal promoter in the expression prediction task naturally led to the question of which sequence motifs within this region were responsible for quantitatively defining expression level. We therefore devised a strategy to identify *k*-mers enriched in the genes predicted to be highly expressed (**Figure 6B**). The distribution of predicted gene expression levels was largely bimodal, allowing the partitioning of genes broadly into genes predicted to have low expression (class A) and high expression (class B). We extracted sub-sequences from the promoters of class B whose saliency scores were higher than the 99th percentile of those observed at the same positions in the promoters of class A. To identify enriched *k*-mers in this set of sub-sequences, we devised a equivalently sized negative set of sub-sequences by permuting the extracted positions in class B to control for positional sequence biases. We then used DREME^52^ to identify *k*-mers enriched in our positive set relative to our negative set. Evaluating the E-values of the top five significantly enriched *k*-mers, we observed the dinucleotide CpG as enriched by orders of magnitude over the second best *k*-mer in both human and mouse species (**Figure 6B**), implicating it as a dominant factor discriminating highly expressed genes from lowly expressed ones. We repeated our procedure on genes in the top half of class B relative to its lower half, and identified an even stronger enrichment of CpG dinucleotides (data not shown). These observations suggest that both the mouse and human models predominantly utilize the spatial distribution of CpG dinucleotides surrounding the proximal promoter to predict the entire continuum of gene expression levels.

The model therefore arrives at a specific prediction: CpG dinucleotides are more enriched in the proximal promoters of highly expressed genes relative to lowly expressed genes. To test this hypothesis, we evaluated the positional enrichment of all 16 possible dinucleotides around the TSS of genes in different gene expression bins relative to chance expectation. While CpGs are globally depleted, genes in higher gene expression bins preserved a greater fraction of CpGs closer to the TSS (**Figure 6C**). This property was true to a much lesser extent for other dinucleotides, with only AA/TT, CA/TG, and CC/GG dinucleotides being able to discriminate between highly and lowly expressed genes in both human and mouse genomes (**Figure S7, Figure S8**).

## DISCUSSION

In this study, we demonstrate that a substantial proportion of variability across genes with respect to their steady-state mRNA expression levels is predictable from features derived solely from genomic sequence. In doing so, our work illustrates—as is the case for gene prediction—that the mathematical function linking genomic sequence to mRNA abundance is in a large part learnable *without* the use of additional sources of experimental data such as those derived from DNase hypersensitivity, TF ChIP, histone ChIP, or MPRAs. Consistent with dogma and recent experiments^23^, we find that the instructions governing transcriptional output are heavily enriched in a gene’s proximal promoter (more specifically, the ±500bp around the TSS). We establish Xpresso as an early initial attempt to confront the problem of gene expression prediction from genomic sequence alone, and anticipate that future algorithms can utilize our effort as a baseline model to improve upon at this prediction task. As deep learning approaches become increasingly sophisticated, we predict that new methods will be able to extract additional spatial relationships and long-range dependencies among motifs that are currently outside of the scope of those learnable by convolutional neural network architectures. While this manuscript was in preparation, another paper was published that similarly demonstrates the feasibility of predicting expression levels from genomic sequence^53^.

Our study provides a theoretical framework to further understand the fundamental question of how different modes of gene regulation contribute to steady-state abundance of mRNA. Querying the performance of the model while considering subsets of features associated with various mechanisms of gene regulation (e.g., mRNA decay and transcription rate) helped dissect their relative contributions to steady-state mRNA levels. Based on the proportion of variance explained by our model thus far, we estimate that between 57-89% of variability can be explained by transcriptional regulation, with the remaining explained by regulation of mRNA stability. These estimates are generally consistent with those derived from experimental measurements in mammalian cells^2,22^. Additionally, we estimate that promoter sequences alone explain ~50% of gene expression variability in humans, again surprisingly consistent with experimentally derived estimates^23^. Collectively, these results reveal that *in silico* strategies to estimate the relative influence of various modes of gene regulation can approximate those more directly relying on experimental measurement.

While our model makes substantial headway into predicting expression levels, between 40-60% of variability still remains unexplained, depending upon the cell type and species considered. We propose that the limitations of our model are also interesting in that they have the potential to inform and resolve the many layers of gene regulation that the model fails to capture. A residual analysis of highly expressed genes that the model under-predicts confirms that enhancers and super-enhancers play a measurably significant role in governing transcriptional programmes. Incorporation of the effects of enhancers in the model is complicated by the difficulty of predicting which promoter(s) any given enhancer influences, as these can be positioned hundreds of kilobases away and skip over genes, as well as the extent to which parameters such as distance modulate the level of enhancer-mediated activation. Such long-range dependencies are poorly modeled by convolutional neural networks. Although the incorporation of distal enhancers into the model has proved to be evasively difficult, the model can be used as a hypothesis generation engine to uncover additional gene regulatory mechanisms that further explain outliers. Our model provides a natural strategy to quantitatively rank candidate silenced and activated genes in different cell types in a way that prioritizes those that most heavily deviate from its predictions. We propose these rankings as a foundation to guide experimentalists interested in dissecting the layers of gene regulation that operate in their cell type of interest.

Scanning large regions of the genome with our pipeline revealed a striking association between regions of high predicted transcriptional activity and CpG islands. While CpG islands are a well-established feature of mammalian genomes that frequently demarcate promoter sequences^49^, our results support the idea that the spatial positioning of CpGs around the proximal promoter is intimately associated with gene expression levels. Our data therefore reinforce the findings from MPRAs that promoter regions enriched with CpG dinucleotides are functionally associated with increased gene expression levels^23^.

Looking forward, we envision the delineation of set of mathematical functions for each cell type that can accurately predict their mRNA expression levels from genome sequence alone as a grand challenge for the field. As shown here, this framework will allow us to quantify and characterize the mechanisms of gene regulation of which we are aware, and may draw our attention to ones that have yet to be discovered.

## METHODS

### Gene expression data collection and pre-processing

We retreived a matrix of normalized expression values for protein-coding mRNAs across 56 tissues and cell lines from RNA-seq data gathered and quantified by the Epigenomics Roadmap Consortium (http://egg2.wustl.edu/roadmap/data/byDataType/rna/expression/57epigenomes.RPKM.pc.gz)^30^.

For mouse gene expression data, we gathered all ENCODE RNA-seq datasets satisfying the following constraints: i) datasets corresponded to “polyA-selected mRNA RNA-seq” or “total RNA-seq”, ii) reads were mapped to the *Mus musculus* mm10 genome assembly, iii) files were “tsv” files corresponding to gene-level quantifications, iv) biosamples were not treated with “DMS”, “LPS”, or “β-estradiol”, v) files were not derived from samples with “low replicate concordance”, “low read depth”, “insufficient read depth”, or “insufficient read length”. Only samples corresponding to the first replicate of each tissue or cell line were utilized. In total, 254 mouse RNA-seq datasets passed these criteria. For cell type-specific questions in the mouse, we used gene expression data from mESCs that were computed previously^15^.

For each species, we computed the median expression level across all cell types for each gene, and transformed all gene expression values 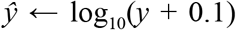 to reduce the right skew of the data. Quantile normalizing the samples of the mouse gene expression matrix prior to computing the median values resulted in nearly an identical set of median expression levels (i.e., 0.9996 correlation before and after quantile normalization), making this step optional.

One-to-one human-to-mouse orthologs were acquired from the Ensembl v90 BioMart^54^ by extracting the “Mouse gene stable ID” and “Mouse homology type” with respect to each human gene.

### Collecting promoter sequences and mRNA half-life features

Human and mouse promoter CAGE peak annotations were downloaded from the FANTOM5 consortium’s UCSC data hub (http://fantom.gsc.riken.jp/5/datahub/hg38/peaks/hg38.cage_peak.bb, hg38 genome build; http://fantom.gsc.riken.jp/5/datahub/mm10/peaks/mm10.cage_peak.bb, mm10 genome build)^47,48^. The best peak corresponding to each promoter, labeled with the keyword “p1@”, was extracted. HUGO gene names or gene name synonyms were converted into Ensembl IDs using the Ensembl v90 BioMart, HGNC ID tables (https://www.genenames.org/cgi-bin/download), and Mouse Genome Informatics gene model tables (http://www.informatics.jax.org/downloads/reports/MGI_Gene_Model_Coord.rpt).

Gene annotations for protein coding genes were derived from Ensembl v90 (hg38 genome build)^54^. Only protein-coding genes were carried forward for analysis, with the following genes filtered out as sources of bias: i) genes located on chrY, whose gene expression depended upon whether their cells of origin were male or female, ii) histone genes, whose expression was mis-quantified due to their mRNAs lacking poly(A) tails, therefore being undersampled in poly(A)-selected RNA-seq libraries. Out of all transcripts corresponding to each gene, the one with the longest ORF, followed by the longest 5′ UTR, followed by the longest 3′ UTR was chosen as the representative transcript for that gene. The G/C content and lengths of each of these functional regions (i.e., 5′ UTRs, ORFs, and 3′ UTRs), intron length, and ORF exon junction density (computed as the number of exon junctions per kilobase of ORF sequence) were gathered as additional features associated with mRNA half-life^21,22^. All length-related features were transformed such that: 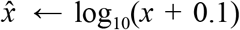 to reduce the right skew, and along with gene expression levels were then z-score normalized by subtracting their respective mean values and dividing by their standard deviations. The starting coordinate of the first exon of the representative transcript was defined as that gene’s transcriptional start site (TSS). The vast majority of mRNAs possessed a dominant CAGE peak; if this was so, the TSS was re-centered to the coordinates of the CAGE peak. While considering CAGE data helped to improve our model, its use was optional as the performance of our models was only slightly worse when considering only Ensembl TSS annotations (r^2^ of 0.54 instead of 0.59 in the human). The ±10 kilobase sequence centered at the TSS was extracted as the putative promoter region to consider. Intermediate steps such as extracting sequences from the genome or converting between bed formats were executed with BEDTools^55^.

### Hyperparameter optimization and model training

Matching gene expression levels to promoter sequences resulted in a total of 18,377 and 21,856 genes in human and mouse, respectively. All continuous variables were mean-centered and scaled to have unit variance, and promoter sequences were one-hot encoded into a boolean matrix. For each species, genes were then randomly partitioned into training, validation, and test sets such that the validation and test sets were allotted 1,000 genes each. We defined the objective function as the minimum mean squared error achieved on the validation set across 10 epochs of training. For the best set of hyperparameters specifying the neural network structure, we trained ten independent trials, and selected the parameters derived from the specific trial and epoch that minimized the validation MSE as our final model. For each trial, parameters were first randomly initialized by sampling from a Glorot normal distribution^56^. Next, the Adam optimizer was used to search for a local minima^56^. The search was performed for 100 epochs but cancelled if a lower validation MSE was not discovered within 7 epochs of the best model discovered so far for that trial. If a cross-validation strategy was implemented, we performed an identical strategy for each of the 10 folds of the data using the respective training and validation sets of the fold. The following software packages were required for model training and testing: Keras 2.0.8^56^, TensorFlow 1.3.0^57^, CUDA 8.0.61, cuDNN 5.1.10, and the Anaconda2 distribution of Python. We initialized a hyperparameter search space to specify the model architecture (**Table 1**), and used Hyperopt^58^ to search for an optimal set of hyperparameters. All models were trained on an NVIDIA Quadro P6000 GPU equipped with 24Gb of video RAM.

Note to reviewers: We intend to release a fully reproducible implementation of the above-described procedures in our Xpresso Github package during the review process.

### Whole-transcriptome predictions

To predict expression levels for all annotated genes, we implemented a 10-fold cross-validation procedure. Specifically, we partitioned the dataset into 10 equally sized bins. For each fold, we i) reserved 1/10 of the data as a test set, 1000 genes as a validation set, and the remaining genes as a training set; ii) trained 10 independent models until convergence, iii) selected the model with the minimum validation mean squared error, and iv) generated a prediction on the test set. We then concatenated all of the predictions together that were derived from the best model from each of the ten folds of the data.

### SuRE MPRA data

Pre-processed genome-wide, stranded SuRE MPRA data mapped to hg19 was acquired as bigwig files from GEO record GSE78709^23^. To compute SuRE activity at a specified promoter, we extracted the mean SuRE signal over covered bases on the correct strand as the gene, centered at 1000bp around the TSS, using utilities provided in the UCSC genome browser (“bigWigAverageOverBed-sampleAroundCenter=1000”)^59^. For the TSS annotations, we utilized our set of CAGE-corrected TSSs lifted over from hg38 to hg19 using liftOver^59^, supplemented with TSSs for genes annotated in Gencode release 27/Ensembl v90 for any missing IDs (https://www.gencodegenes.org/releases/27lift37.html)^60^.

### Baseline models

To train baseline models for comparison (**Figure S4**), we merged the training and validation sets used initially for hyperparameter optimization and model training. For each gene, we first extracted the ±1500bp window centered each TSS and defined this as the promoter. For *k*-mer-based models, we counted the frequency of all *k*-mers occuring occurring in the promoter region, varying *k* from 1 to 5 and 6 for human and mouse, respectively. To train transcription-factor-based models, we scanned promoters using FIMO^61^ using positional weight matrices derived from the JASPAR 2016 Core Vertebrate set^62^. Default parameters were used for the search, except that the set of promoter sequences to compute a first order Markov background model for the search. For the transcription factors matched to the promoters, we populated a binary matrix with a 1 if a significant motif was detected for the promoter, and 0 otherwise. We then trained multiple linear regression models explaining median mRNA expression levels as a function of i) the collection of *k*-mer counts, ii) the binary matrix of JASPAR matches, or iii) both of the former. These models were trained both with and without half-life features, with the r^2^ evaluated on the test set.

### Prediction on a genomic window

We extracted 10.5Kb sequences tiling across the 600 Kb and 700 Kb regions of the human and mouse genomes, respectively, in 100bp increments (Genome coordinates: chr1:109500000-110300000, hg19 genome build; chr3:107800000-108500000, mm10 genome build). The “bedtools makewindows” (parameters “-w 10500 -s 100”) was used to generate these windows, and “bedtools getfasta” to extract the sequences55. Predictions were then made using our cell type-agnostic Xpresso model trained upon median gene expression data, using zero values for all half-life features (with zero corresponding to the mean values as these features had been z-score normalized).

---

**Figure S1.**
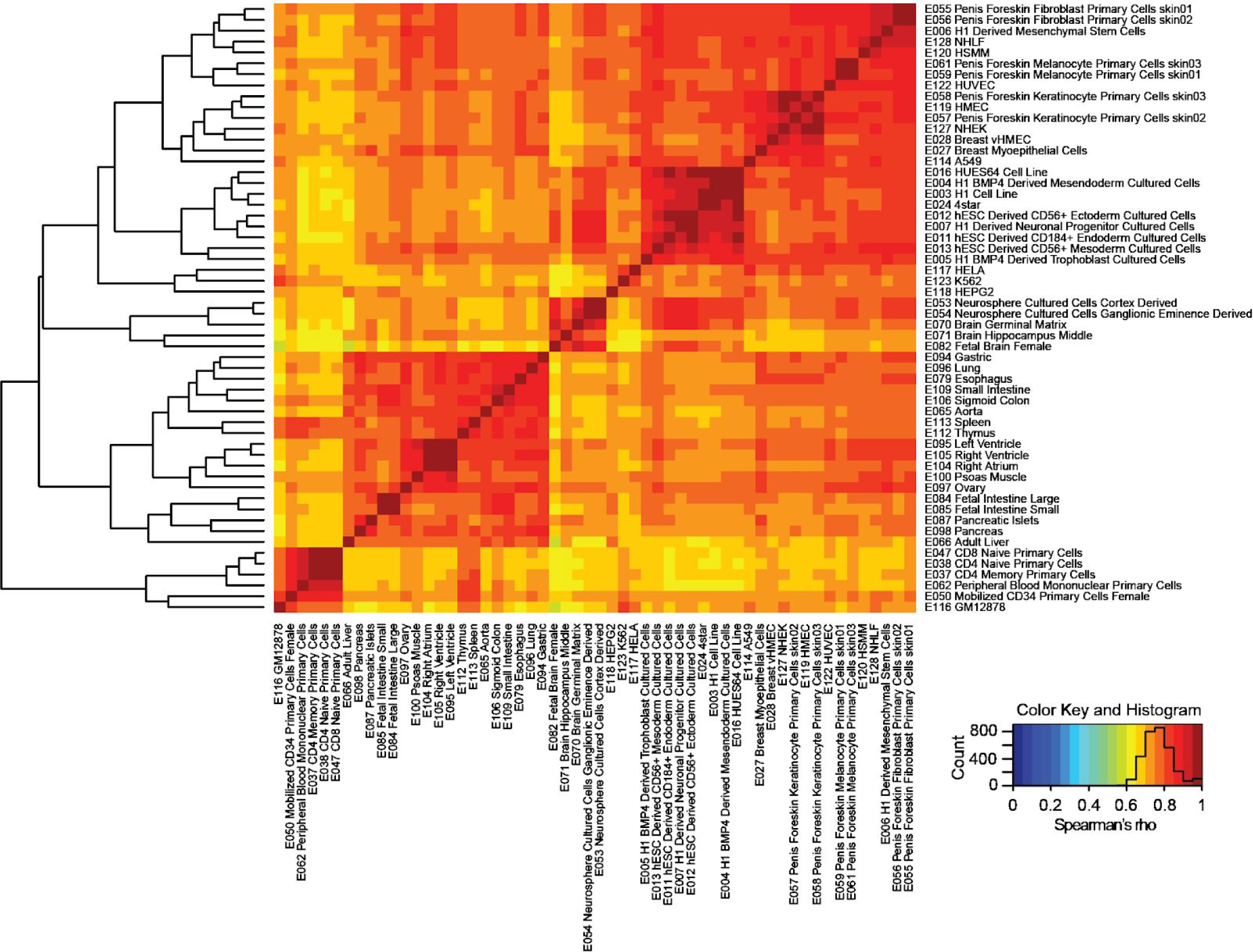
Gene expression levels are highly concordant across cell types. Heatmap of the pairwise Spearman correlation values measured between mRNA expression levels derived from each pair of cell types from 56 cell types aggregated by the Epigenomics Roadmap Consortium^30^, clustered using hierarchical clustering according to the indicated dendrogram. Labeled is the 4-letter Epigenomics Roadmap code alongside the corresponding cell type (e.g. E123 indicates K562 cells). Overlayed in the color key is a histogram of all correlation values.

**Figure S2.**
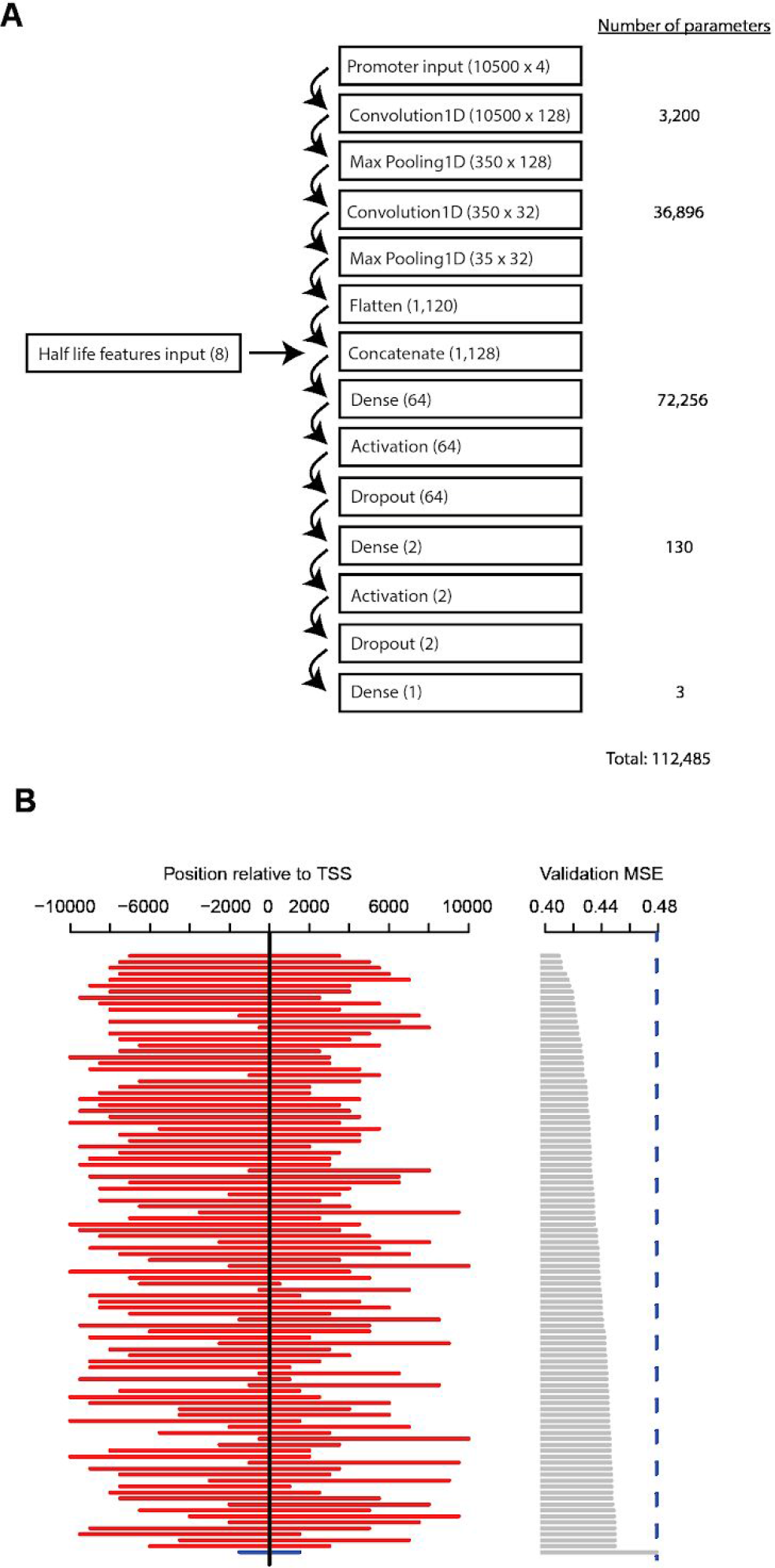
Properties of deep learning architecture learned during hyperparameter search. **A)** Structure of the best hyperparameter architecture uncovered. Within each box is the indicated type of Keras layer as well as the size of the vector or matrix associated with the layer for a single training example, indicated in parenthesis. Specified to the right of the flow diagram are the number of free parameters fit for each layer. **B)** Optimal and suboptimal promoter windows discovered during hyperparameter search as well as the best validation mean squared error (MSE) corresponding to given hyperparameter configuration, ranked according to MSE. Labeled in blue is the window associated with the best manually discovered hyperparameter configuration, with the blue dashed line indicating the associated validation MSE for the specified architecture.

**Figure S3.**
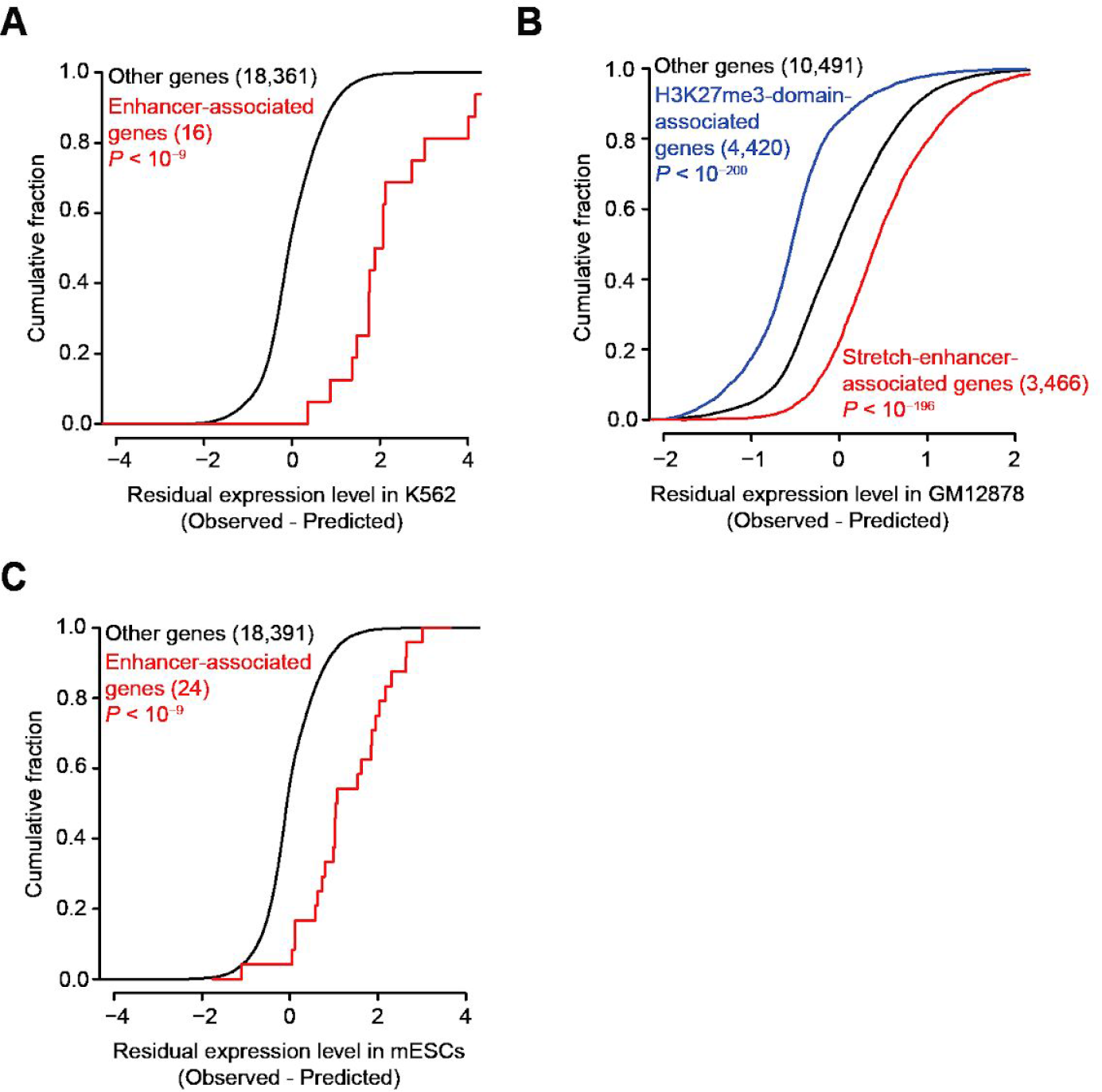
Additional gene regulatory mechanisms associated with the residuals of cell type-specific models. **A)** This panel is similar to **Figure 3B**, except that it displays cumulative distributions of residuals corresponding to the enhancer-associated genes shown in **Figure 3A**. **B)** This panel is similar to **Figure 3B**, except that it evaluates the relationships of stretch-enhancer-associated genes and H3K27me3-domain-associated genes in GM12878 cells. **C)** This panel is similar to **Figure 3E**, except that it displays cumulative distributions of residuals corresponding to the enhancer and super-enhancer-associated genes shown in **Figure 3D**.

**Figure S4.**
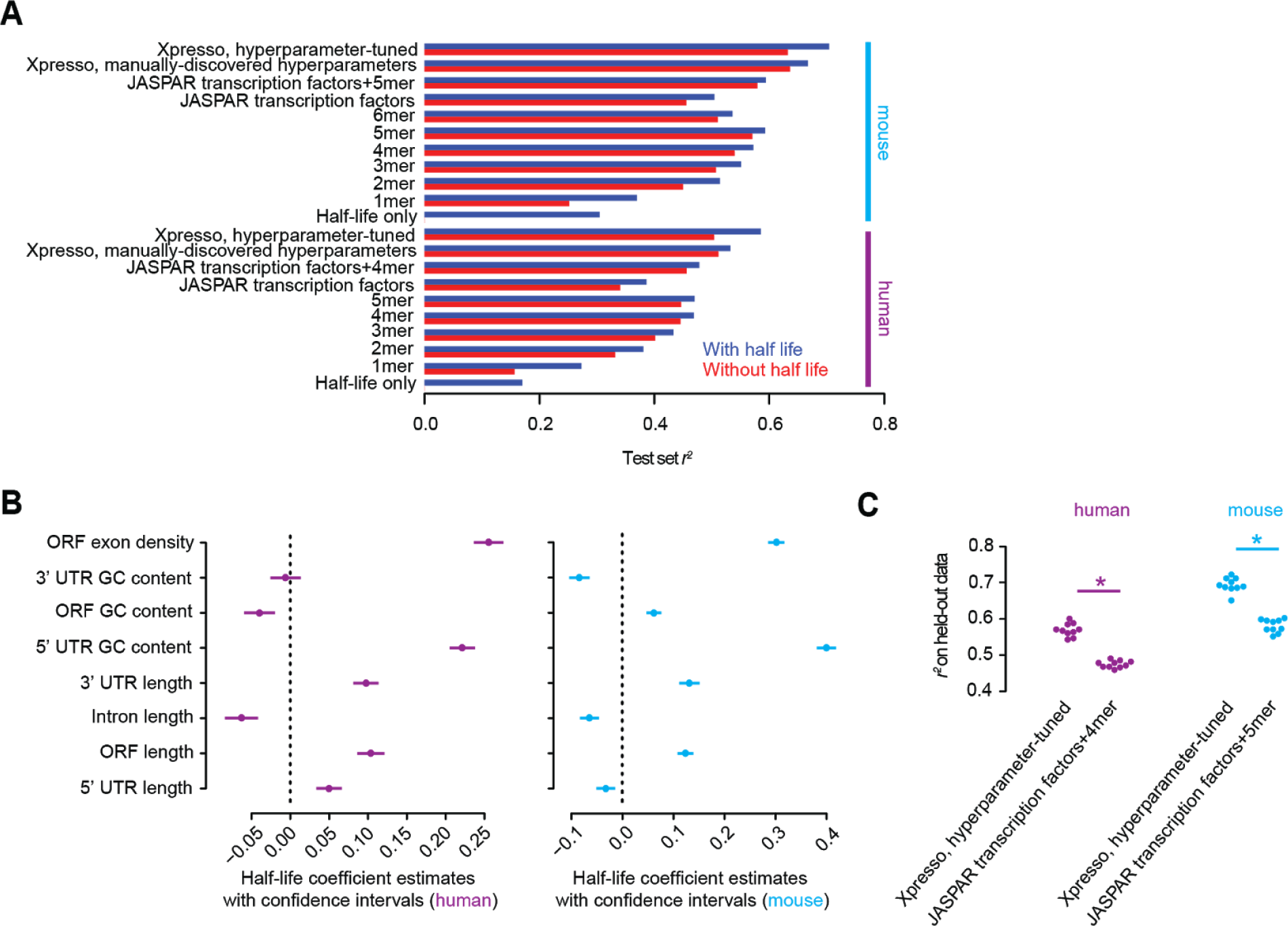
Predictive performance of baseline models. **A)** Comparison of deep learning-based models relative to several baseline models in explaining the variance in median mRNA expression levels on a test set. All human and mouse models were trained with and without half-life information. Baseline models were trained using multiple linear regression, using the following features as input: i) *k*-mer counts in a ±1500bp window centered each TSS, varying *k* from 1 to 5 or 6 depending upon the optimal *k* at which performance plateaued, ii) counts of significant hits derived from motif scans of JASPAR transcription factor sites in the same window, or iii) both counts from motif scans of JASPAR transcription factor sites and *k*-mer counts in the same window. **B)** Coefficients corresponding to each individual feature from the “half-life only” human and mouse models in panel (A). Shown is the mean and 95% confidence intervals for each feature. **C)** Comparison of 10-fold cross-validated r2 between the best hyperparameter-tuned Xpresso model and the best alternative sequence-based model shown in panel (A), for both human and mouse species. Shown are the 10 individual r2 values for each fold of the held-out data in a beeswarm plot. Significance was computed using a paired two-sample t-test (\**P* < 10^−8^).

**Figure S5.**
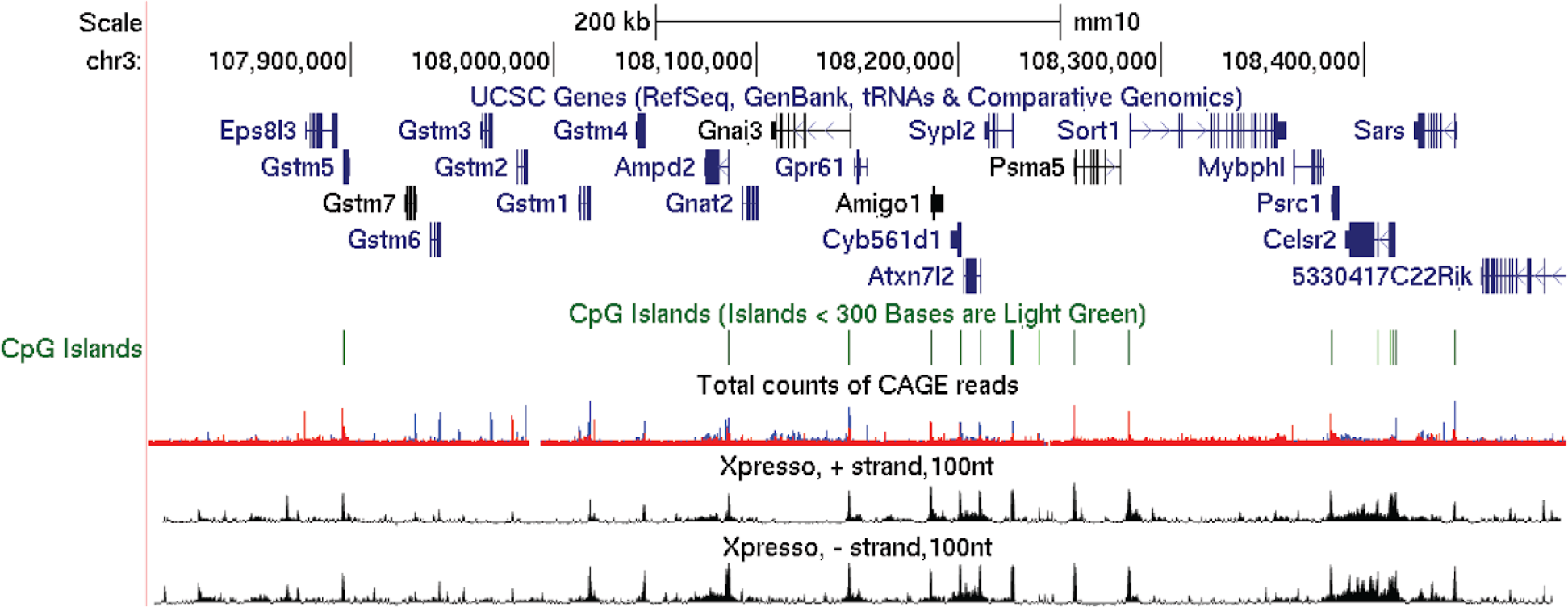
Xpresso predictions recapitulate transcriptional activity across a 700Kb region of mouse chromosome 3. Shown are the predicted Xpresso-predicted expression levels in 100-nt increments from the plus and minus strands of genomic sequence of the mouse mm10 genome assembly. Also shown are gene annotations, CpG island calls, and genome-wide CAGE signal aggregated among many cell types (with red indicating signal from the positive strand and blue indicating signal from the negative strand). See also **Figure 5** for predictions on the syntenic human locus.

**Figure S6.**
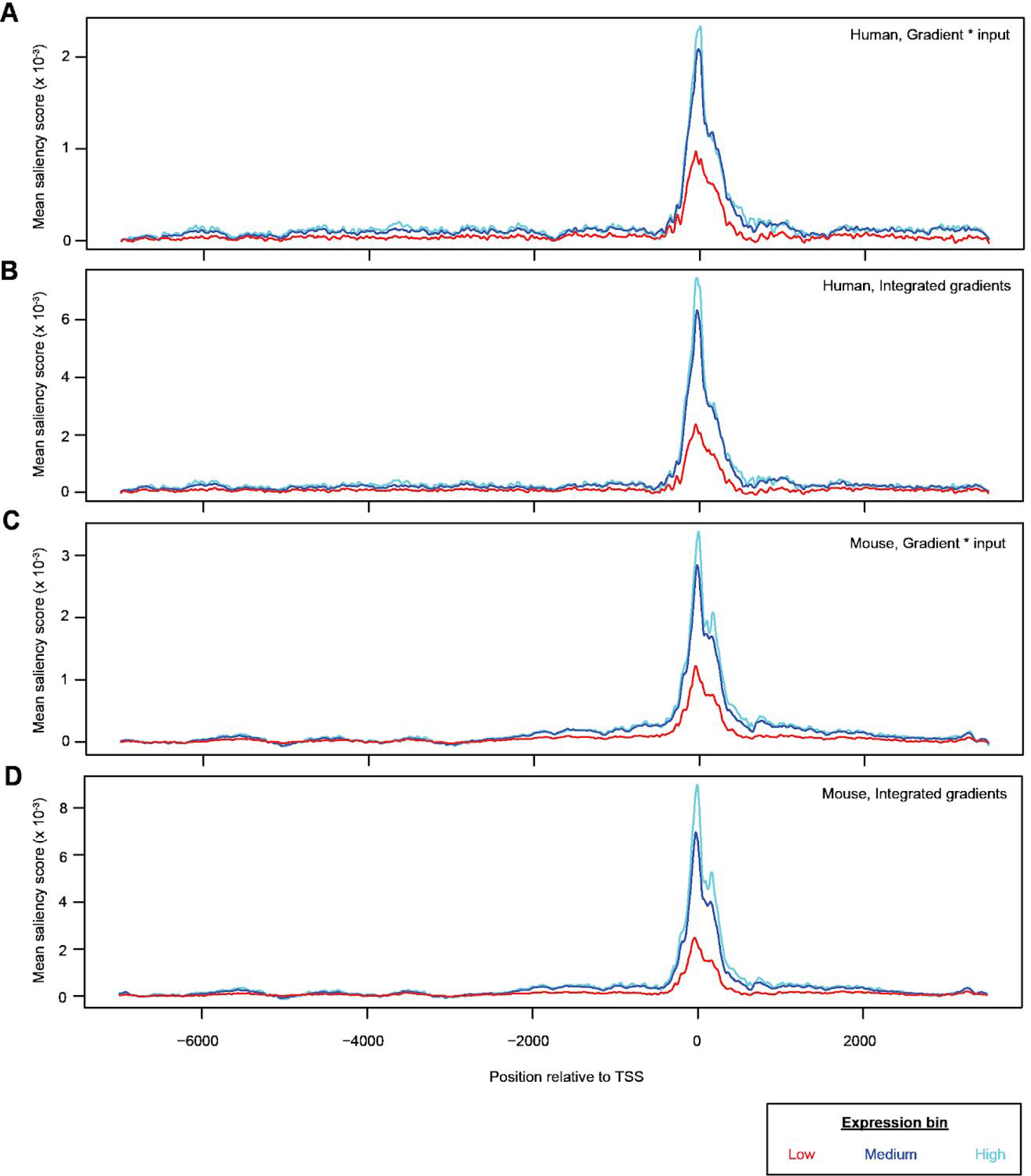
Xpresso isolates the mammalian proximal promoter as the key region relevant for the prediction of gene expression. **A)** This panel is identical to that of **Figure 6A** except that it shows saliency scores from the entire −7000 to +3500 region of the input window surrounding the TSS for human promoters. **B)** This panel is similar to those of panel (A) except that the “Integrated gradients” approach is used instead of the “Gradient * input” approach. **C-D)** These panels are similar to those of panels (A-B) except that they show the results using mouse promoters.

**Figure S7.**
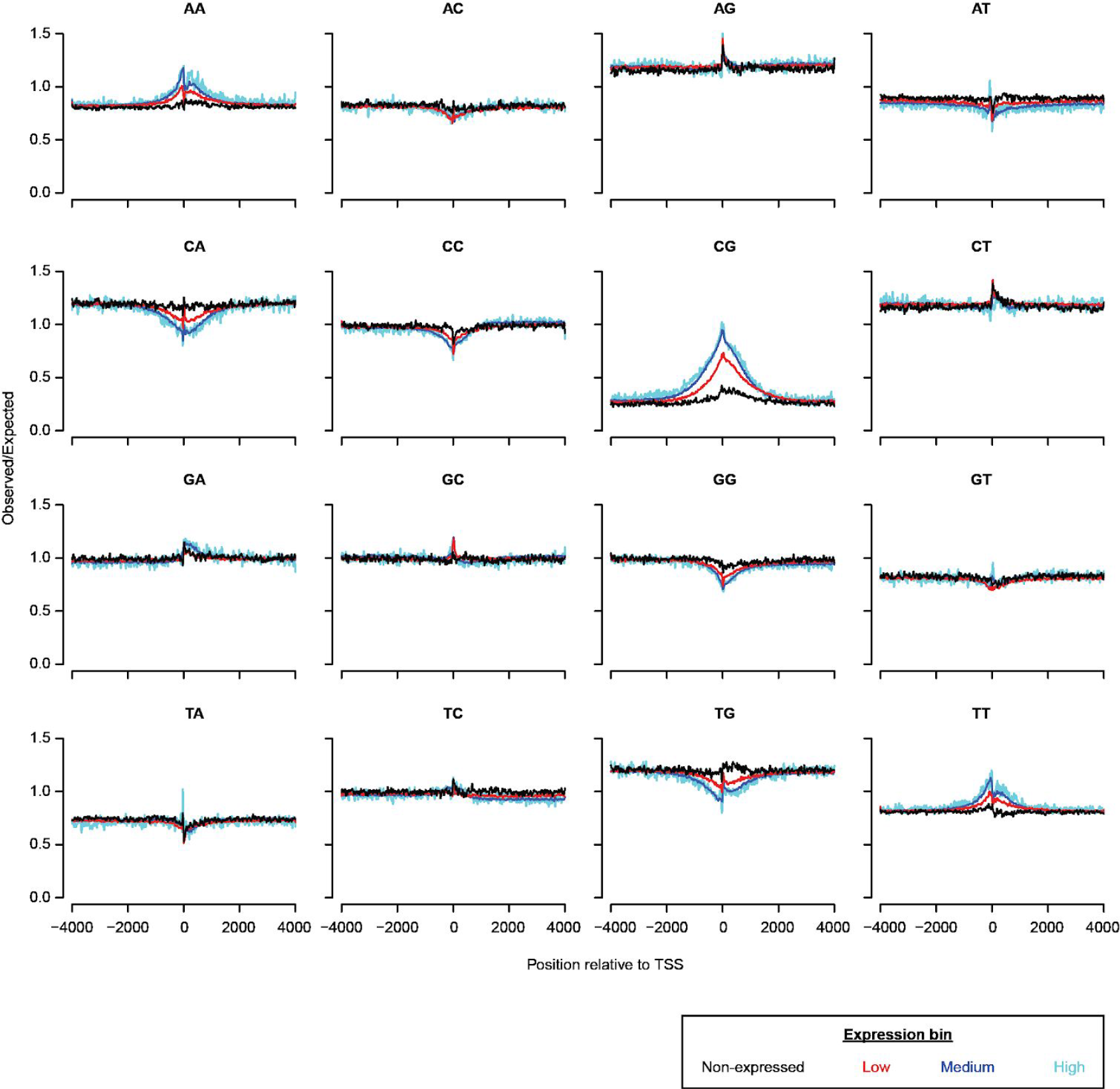
Relationship between dinucleotide content around the TSS and gene expression levels in the human. This panel is similar to that of **Figure 6C**. Positional enrichment of dinucleotides relative to chance expectation in human promoters for different gene expression level bins. The expected frequency of dinucleotides is computed as the joint probability of the composite mononucleotides co-occurring based upon their frequency in the corresponding position and gene expression bin.

**Figure S8.**
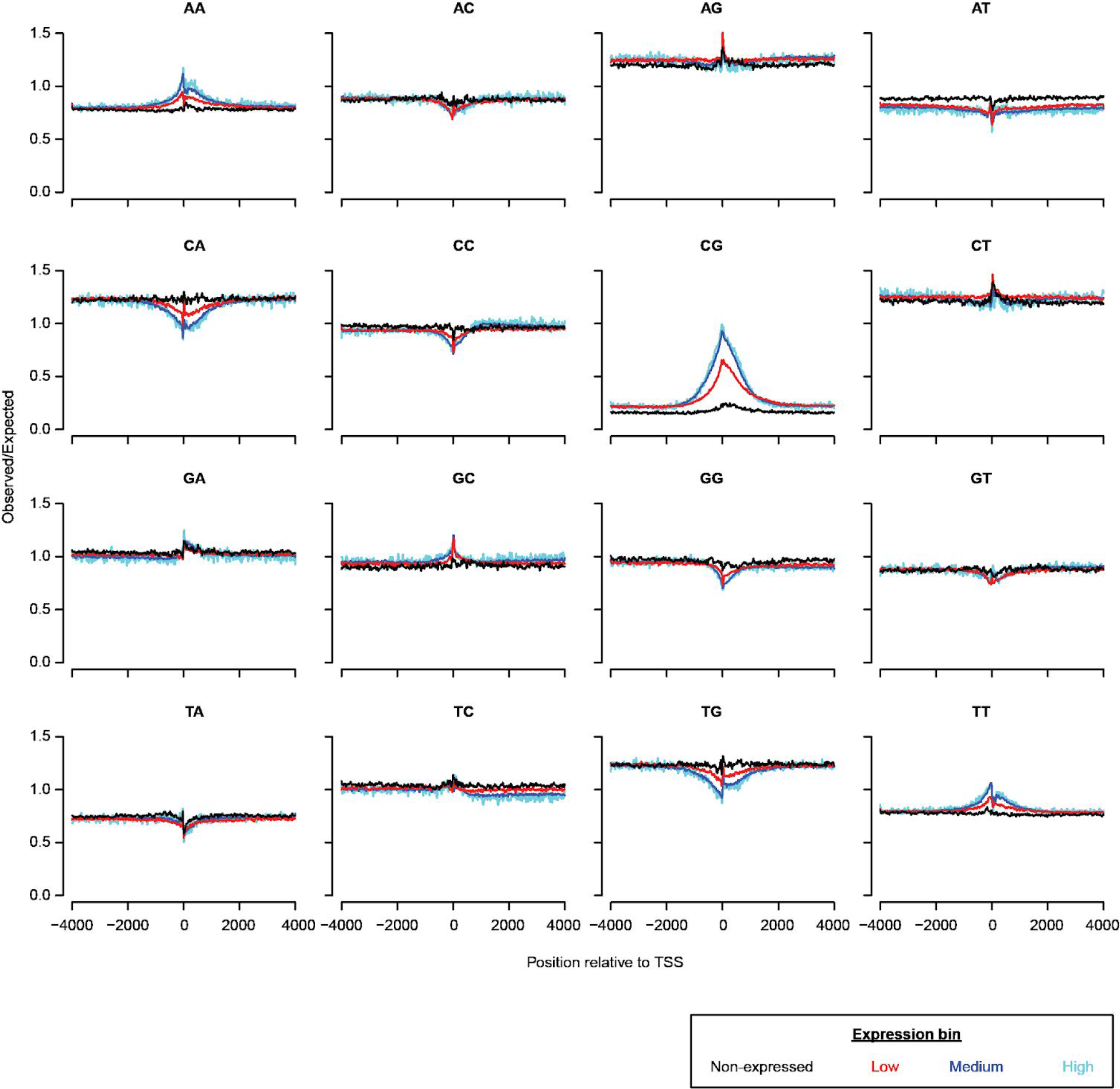
Relationship between dinucleotide content around the TSS and gene expression levels in the mouse. This panel is similar to that of **Figure S7**, except it shows information for mouse promoters instead of human promoters.

## DATA ACCESS

Associated code necessary to train and test the deep learning models and reproduce each figure of the manuscript will be made available upon publication through a GitHub package.

## ACKNOWLEDGMENTS

We thank J. Tome, S. Kim, and other members of the Shendure lab for critical commentary and helpful discussions. This material is based upon work supported under an NRSA NIH fellowship 5T32HL007093 (to V.A.) and NIH grants 5DP1HG007811, 5R01HG009136, and 5R01CA197139 (to J.S.). J.S. is an investigator of the Howard Hughes Medical Institute.

## REFERENCES CITED

1 Schwanhausser, B. et al. Global quantification of mammalian gene expression control. Nature 473, 337–342 (2011).

2 Li, J. J., Bickel, P. J. & Biggin, M. D. System wide analyses have underestimated protein abundances and the importance of transcription in mammals. PeerJ 2, e270 (2014).

3 Vogel, C. et al. Sequence signatures and mRNA concentration can explain two-thirds of protein abundance variation in a human cell line. Mol. Syst. Biol. 6, 400 (2010).

4 Edfors, F. et al. Gene-specific correlation of RNA and protein levels in human cells and tissues. Mol. Syst. Biol. 12, 883 (2016).

5 Cao, R. et al. Role of histone H3 lysine 27 methylation in Polycomb-group silencing. Science 298, 1039–1043 (2002).

6 Parker, S. C. et al. Chromatin stretch enhancer states drive cell-specific gene regulation and harbor human disease risk variants. Proc. Natl. Acad. Sci. U. S. A. 110, 17921–17926 (2013).

7 Whyte, W. A. et al. Master transcription factors and mediator establish super-enhancers at key cell identity genes. Cell 153, 307–319 (2013).

8 Agarwal, V., Bell, G. W., Nam, J. W. & Bartel, D. P. Predicting effective microRNA target sites in mammalian mRNAs. Elife 4, (2015).

9 Wickens, M., Bernstein, D. S., Kimble, J. & Parker, R. A PUF family portrait: 3 ’ UTR regulation as a way of life. Trends Genet. 18, 150–157 (2002).

10 Brennan, C. M. & Steitz, J. A. HuR and mRNA stability. Cell. Mol. Life Sci. 58, 266–277 (2001).

11 Cheng, C. et al. A statistical framework for modeling gene expression using chromatin features and application to modENCODE datasets. Genome Biol. 12, (2011).

12 Dong, X. J. et al. Modeling gene expression using chromatin features in various cellular contexts. Genome Biol. 13, (2012).

13 Cheng, C. et al. Understanding transcriptional regulation by integrative analysis of transcription factor binding data. Genome Res. 22, 1658–1667 (2012).

14 McLeay, R. C., Lesluyes, T., Cuellar Partida, G. & Bailey, T. L. Genome-wide in silico prediction of gene expression. Bioinformatics 28, 2789–2796 (2012).

15 Ouyang, Z., Zhou, Q. & Wong, W. H. ChIP-Seq of transcription factors predicts absolute and differential gene expression in embryonic stem cells. Proc. Natl. Acad. Sci. U. S. A. 106, 21521–21526 (2009).

16 Schmidt, F. et al. Combining transcription factor binding affinities with open-chromatin data for accurate gene expression prediction. Nucleic Acids Res. 45, 54–66 (2017).

17 Karlic, R., Chung, H. R., Lasserre, J., Vlahovicek, K. & Vingron, M. Histone modification levels are predictive for gene expression. Proc. Natl. Acad. Sci. U. S. A. 107, 2926–2931 (2010).

18 Jain, D., Baldi, S., Zabel, A., Straub, T. & Becker, P. B. Active promoters give rise to false positive ‘Phantom Peaks’ in ChIP-seq experiments. Nucleic Acids Res. 43, 6959–6968 (2015).

19 Teytelman, L., Thurtle, D. M., Rine, J. & van Oudenaarden, A. Highly expressed loci are vulnerable to misleading ChIP localization of multiple unrelated proteins. Proc. Natl. Acad. Sci. U. S. A. 110, 18602–18607 (2013).

20 Krebs, W. et al. Optimization of transcription factor binding map accuracy utilizing knockout-mouse models. Nucleic Acids Res. 42, 13051–13060 (2014).

21 Sharova, L. V. et al. Database for mRNA half-life of 19 977 genes obtained by DNA microarray analysis of pluripotent and differentiating mouse embryonic stem cells. DNA Res. 16, 45–58 (2009).

22 Spies, N., Burge, C. B. & Bartel, D. P. 3’ UTR-isoform choice has limited influence on the stability and translational efficiency of most mRNAs in mouse fibroblasts. Genome Res. 23, 2078–2090 (2013).

23 van Arensbergen, J. et al. Genome-wide mapping of autonomous promoter activity in human cells. Nat. Biotechnol. (2016). doi:10.1038/nbt.3754

24 Segal, E. & Widom, J. From DNA sequence to transcriptional behaviour: a quantitative approach. Nat. Rev. Genet. 10, 443–456 (2009).

25 Angermueller, C., Parnamaa, T., Parts, L. & Stegle, O. Deep learning for computational biology. Mol. Syst. Biol. 12, 878 (2016).

26 Alipanahi, B., Delong, A., Weirauch, M. T. & Frey, B. J. Predicting the sequence specificities of DNA- and RNA-binding proteins by deep learning. Nat. Biotechnol. 33, 831–838 (2015).

27 Zhou, J. & Troyanskaya, O. G. Predicting effects of noncoding variants with deep learning-based sequence model. Nat. Methods 12, 931–934 (2015).

28 Kelley, D. R., Snoek, J. & Rinn, J. L. Basset: learning the regulatory code of the accessible genome with deep convolutional neural networks. Genome Res. 26, 990–999 (2016).

29 Kelley, D. R. et al. Sequential regulatory activity prediction across chromosomes with convolutional neural networks. Genome Res. 161851 (2018).

30 Roadmap Epigenomics, Consortium et al. Integrative analysis of 111 reference human epigenomes. Nature 518, 317–330 (2015).

31 Bergstra, J. S., Bardenet, R., Bengio, Y. & Kégl, B. Algorithms for hyper-parameter optimization. (2011).

32 Fulco, C. P. et al. Systematic mapping of functional enhancer-promoter connections with CRISPR interference. Science 354, 769–773 (2016).

33 Klann, T. S. et al. CRISPR-Cas9 epigenome editing enables high-throughput screening for functional regulatory elements in the human genome. Nat. Biotechnol. 35, 561–568 (2017).

34 Xie, S., Duan, J., Li, B., Zhou, P. & Hon, G. C. Multiplexed Engineering and Analysis of Combinatorial Enhancer Activity in Single Cells. Mol. Cell 66, 285–299 e5 (2017).

35 Marco, E. et al. Multi-scale chromatin state annotation using a hierarchical hidden Markov model. Nat. Commun. 8, 15011 (2017).

36 Schofield, J. A., Duffy, E. E., Kiefer, L., Sullivan, M. C. & Simon, M. D. TimeLapse-seq: adding a temporal dimension to RNA sequencing through nucleoside recoding. Nat. Methods 15, 221–225 (2018).

37 Moorthy, S. D. et al. Enhancers and super-enhancers have an equivalent regulatory role in embryonic stem cells through regulation of single or multiple genes. Genome Res. 27, 246–258 (2017).

38 Boyer, L. A. et al. Polycomb complexes repress developmental regulators in murine embryonic stem cells. Nature 441, 349–353 (2006).

39 Herzog, V. A. et al. Thiol-linked alkylation of RNA to assess expression dynamics. Nat. Methods 14,1198–1204 (2017).

40 Denzler, R. et al. Impact of MicroRNA Levels, Target-Site Complementarity, and Cooperativity on Competing Endogenous RNA-Regulated Gene Expression. Mol. Cell (2016). doi:10.1016/j.molcel.2016.09.027

41 Mullokandov, G. et al. High-throughput assessment of microRNA activity and function using microRNA sensor and decoy libraries. Nat. Methods 9, 840–846 (2012).

42 Abdalla, M., Abdalla, M., McCarthy, M. I. & Holmes, C. C. A general framework for predicting the transcriptomic consequences of non-coding variation. bioRxiv 279323 (2018). doi:10.1101/279323

43 Cooper, S. J., Trinklein, N. D., Anton, E. D., Nguyen, L. & Myers, R. M. Comprehensive analysis of transcriptional promoter structure and function in 1% of the human genome. Genome Res. 16, 1–10 (2006).

44 Landolin, J. M. et al. Sequence features that drive human promoter function and tissue specificity. Genome Res. 20, 890–898 (2010).

45 Nguyen, T. A. et al. High-throughput functional comparison of promoter and enhancer activities. Genome Res. 26, 1023–1033 (2016).

46 Duren, Z., Chen, X., Jiang, R., Wang, Y. & Wong, W. H. Modeling gene regulation from paired expression and chromatin accessibility data. Proc. Natl. Acad. Sci. U. S. A. 114, E4914–E4923 (2017).

47 Lizio, M. et al. Gateways to the FANTOM5 promoter level mammalian expression atlas. Genome Biol. 16, 22 (2015).

48 Fantom Consortium et al. A promoter-level mammalian expression atlas. Nature 507, 462–470 (2014).

49 Gardiner-Garden, M. & Frommer, M. CpG islands in vertebrate genomes. J. Mol. Biol. 196, 261–282 (1987).

50 Ernst, J. & Kellis, M. ChromHMM: automating chromatin-state discovery and characterization. Nat. Methods 9, 215–216 (2012).

51 Ancona, M., Ceolini, E., Oztireli, C. & Gross, M. Towards better understanding of gradient-based attribution methods for Deep Neural Networks. in 6th International Conference on Learning Representations (ICLR 2018) (research-collection.ethz.ch, 2018).

52 Bailey, T. L. DREME: motif discovery in transcription factor ChIP-seq data. Bioinformatics 27,1653–1659 (2011).

53 Zhou, J. et al. Deep learning sequence-based ab initio prediction of variant effects on expression and disease risk. Nat. Genet. 50, 1171–1179 (2018).

54 Aken, B. L. et al. Ensembl 2017. Nucleic Acids Res. 45, D635–D642 (2016).

55 Quinlan, A. R. & Hall, I. M. BEDTools: a flexible suite of utilities for comparing genomic features. Bioinformatics 26, 841–842 (2010).

56 Chollet, F. Keras. (2015).

57 Abadi, M. et al. Tensorflow: Large-scale machine learning on heterogeneous distributed systems. arXiv preprint arXiv:1603.04467 (2016).

58 Bergstra, J., Yamins, D. & Cox, D. Making a science of model search: Hyperparameter optimization in hundreds of dimensions for vision architectures. (2013).

59 Kent, W. J. et al. The human genome browser at UCSC. Genome Res. 12, 996–1006 (2002).

60 Harrow, J. et al. GENCODE: the reference human genome annotation for The ENCODE Project. Genome Res. 22, 1760–1774 (2012).

61 Grant, C. E., Bailey, T. L. & Noble, W. S. FIMO: scanning for occurrences of a given motif. Bioinformatics 27, 1017–1018 (2011).

62 Mathelier, A. et al. JASPAR 2016: a major expansion and update of the open-access database of transcription factor binding profiles. Nucleic Acids Res. 44, D110–D115 (2015).

63 Antequera, F. Structure, function and evolution of CpG island promoters. Cell. Mol. Life Sci. 60,1647–1658 (2003).

